# Network Inference with Granger Causality Ensembles on Single-Cell Transcriptomic Data

**DOI:** 10.1101/534834

**Authors:** Atul Deshpande, Li-Fang Chu, Ron Stewart, Anthony Gitter

**Author notes:** Department of Oncology, Johns Hopkins University, Baltimore, MD 21205, USA. Department of Comparative Biology and Experimental Medicine, University of Calgary, Calgary, AB T2N 4N1, Canada.

## Abstract

Advances in single-cell transcriptomics enable measuring the gene expression of individual cells, allowing cells to be ordered by their state in a dynamic biological process. Many algorithms assign ‘pseudotimes’ to each cell, representing the progress along the biological process. Ordering the expression data according to such pseudotimes can be valuable for understanding the underlying regulator-gene interactions in a biological process, such as differentiation. However, the distribution of cells sampled along a transitional process, and hence that of the pseudotimes assigned to them, is not uniform. This prevents using many standard mathematical methods for analyzing the ordered gene expression states. We present Single-cell Inference of Networks using Granger Ensembles (SINGE), an algorithm for gene regulatory network inference from single-cell gene expression data. Given ordered single-cell data, SINGE uses kernel-based Granger Causality regression, which smooths the irregular pseudotimes and missing expression values. It then aggregates the predictions from an ensemble of regression analyses with a modified Borda count to compile a ranked list of candidate interactions between transcriptional regulators and their target genes. In two mouse embryonic stem cell differentiation case studies, SINGE outperforms other contemporary algorithms for gene network reconstruction. However, a more detailed examination reveals caveats about transcriptional network reconstruction with single-cell RNA-seq data. Network inference methods, including SINGE, may have near random performance for predicting the targets of many individual regulators even if the overall performance is good. In addition, including uninformative pseudotime values can hurt the performance of network reconstruction methods. A MATLAB implementation of SINGE is available at https://github.com/gitter-lab/SINGE.

## 1. Introduction

Identifying the underlying gene regulatory networks (GRNs) that dictate cell-fate decisions is important for understanding biological systems. Although RNA-seq experiments on populations of cells have been used to study cellular decision making, averaging transcriptional information from a heterogeneous population of cells can obscure biological signals. Advances in single-cell transcriptomics, such as single-cell RNA-seq, have enabled observing the gene expression states of individual cells [1–3]. While these solve the averaging problem faced by bulk transcriptomics, they are beset with new technical challenges, including measurement dropouts and a lower signal-to-noise ratio. Despite the technical problems, snapshots of the gene expression states of individual cells provide larger sample sizes and a finer understanding of the gene expression and regulatory dynamics during a biological process.

Many algorithms use single-cell RNA-seq data to infer GRNs [4–6], taking advantage of the large sample sizes. In the strictest sense, GRNs only include regulation of genes by transcription factors (TF). However, we use the term GRN to mean a network of directed causal relationships between any regulators (not necessarily TFs) and their target genes. GRN inference requires identifying relationships between transcriptional regulators and their target genes or gene modules [7–9]. One strategy is to search gene expression datasets for dependencies among mRNA expression levels, making the simplifying assumption that a regulator’s mRNA level approximates its regulatory activity. Single-cell datasets offer more data from which to learn these gene-gene relationships using multivariate information theory [10], linear regression [11], or other approaches. Methods like GENIE3 [12], which were originally designed to infer GRNs from bulk transcriptomic data using tree-based ensembles, can be easily adapted for single-cell datasets. When single-cell expression data are collected at multiple times points, it provides more information that can be used for GRN inference. GRN reconstruction methods originally designed for bulk time-series transcriptomic data [13] can be repurposed to analyze time-stamped single-cell data. For example, Jump3 [14], a hybrid machine learning and model-based approach, has been adapted in this manner [15]. Time-stamped single-cell data also enables analyzing the evolution of gene expression distributions over time [16], which is not possible with bulk time series data or single-cell data collected at one time point.

When single-cell RNA-seq samples are not collected at multiple time points, computationally ordering cells along a biological process based on their expression states can approximate each cell’s position along the process. These inferred times, called ‘pseudotimes’, can potentially lead to greater understanding of the causal regulatory relationships between genes. The dozens of algorithms for ordering cells and assigning pseudotimes [17], also referred to as trajectory inference, can be distinguished by their use of prior knowledge, treatment of pseudotime uncertainty, and the supported trajectory types [18]. Pseudotime algorithms can target cyclic [19, 20], linear [21, 22], bifurcating [23], multifurcating [24], or tree-structured [25, 26] trajectories. In most of these methods, a numeric pseudotime is assigned to each cell, which represents the cell’s progress along the trajectory.

Similar to time series data, the pseudotemporal ordering provides an understanding of the gene expression trends along the biological process, which can support more accurate GRN reconstruction. Strategies for GRN inference with pseudotemporal data are related to those for time-stamped data with additional specializations to account for the technical differences. For example, SINCERITIES [27], originally designed to infer GRNs using Granger Causality-inspired ridge regression on time-stamped expression data, also admits pseudotime-labelled cells. SCODE [15], GRISLI [28], and Ocone et al. [29] infer GRNs by modelling the cell dynamics as ordinary differential equations with pseudotime as the temporal reference. Other strategies involve Gaussian processes regression for smoothing pseudotemporal data [30], time-lagged correlation [31], variational Bayesian inference on a first-order autoregressive moving average model [32], modified Restricted Directed Information [33], unsupervised classification using Gaussian Mixture Models [34], empirical Bayes-based thresholding [35], modeling information propagation through genes as a cascade [36], and transfer entropy [37]. These strategies require estimating the cell trajectories before GRN inference. An alternative approach is to perform joint trajectory and co-expression network inference, for example, using Ornstein-Uhlenbeck models [24] or Gaussian mixtures with continuous parameters [38]. Despite these algorithmic advances, in case studies on real data the GRN reconstruction performance has often been disappointing and sometimes not substantially better than random networks.

In this study, we adapt Granger Causality for pseudotemporally-ordered single-cell expression data to assess whether this causal framework can overcome the difficulties faced by prior pseudotime-based GRN inference methods. We introduce our Single-cell Inference of Networks using Granger Ensembles (SINGE) algorithm, an ensemble-based GRN reconstruction technique that uses modified Granger Causality on single-cell data annotated with pseudotimes. Granger Causality [39, 40] is a powerful approach for detecting specific types of causal relationships in long time series data. It has been used with bulk times series gene expression data [41–46], but these time series are typically short due to experimental limitations, making it more difficult to detect reliable gene-gene dependencies. The longer (pseudo) time series obtained from ordered single-cell datasets make them appealing for Granger Causality-based GRN reconstruction. However, single-cell challenges such as dropouts and irregular sampling along the biological trajectory counteract the benefits of the longer pseudotime series. SINGE addresses these concerns by using a kernel-based Granger Causality method that smooths the expression data and ensembling to improve GRN prediction robustness.

We apply SINGE to reconstruct GRNs of two mouse embryonic stem cell differentiation processes characterized with single-cell RNA-seq. SINGE compares favorably with existing GRN inference methods when evaluated using ChIP-seq, ChIP-chip, loss-of-function, and gain-of-function data. However, our evaluation reveals important caveats about GRN evaluation and the value of pseudotime for GRN inference that are broadly applicable for pseudotime-based GRN reconstruction.

## 2. Results

### 2.1. SINGE and Granger Causality Overview

SINGE takes ordered single-cell gene expression data as input and provides a ranked list of regulator-gene relationships as its primary output. It requires the single-cell dataset to be annotated with pseudotimes. This assigns a numeric pseudotime to each cell in the dataset that represents how much that cell has progressed through a dynamic biological process such as differentiation. For each target gene, SINGE assesses which past expression values are most predictive of its expression, that is, the candidate regulators of each gene. The lagged dependencies are detected using a specialized form of Granger Causality, which is framed as a regularized regression problem. The past expression values are determined using the pseudotimes.

The Granger Causality [39, 40] test at SINGE’s core is a hypothesis test to ascertain predictive causality between a ‘source’ and ‘target’ time series. A series *x* is said to Granger-cause *y* if past values of *x* contain information that helps predict future values of *y*. The primary complication of applying Granger Causality to single-cell expression data with inferred pseudotimes is that the distribution of cells along the trajectory, and the pseudotimes assigned to them, is not uniform. Standard Granger Causality is not an effective analytical tool with irregularly-spaced pseudotimes [33]. One potential workaround is to resample the irregularly-spaced pseudotime series to obtain a regular time series. However, resampling introduces interpolation errors in the form of a low-pass filtering, which could be detrimental to analysis of highly non-linear biological processes. SINGE instead uses an alternative solution proposed by Bahadori and Liu, the Generalized Lasso Granger (GLG) test [47]. GLG modifies the Lasso Granger test [48] to support irregular time series. Within SINGE, GLG uses a kernel function to smooth the past expression values of candidate regulators, mitigating the irregularly-spaced pseudotimes and zero values that are prevalent in single-cell expression data. Our Granger Causality formulation models a strict delay between the regulator and target gene expression in pseudotime. The time-lagged expression relationships between regulators and target genes are motivated by simulations of transcription kinetics that naturally induce such lags [49, 50]. Therefore, SINGE is not intended to identify any ‘instantaneous’ regulatory relationships.

SINGE depends on hyperparameters that control the kernel smoothing, sparsity, and which window of previous expression is considered. We do not search for a single optimal set of hyperparameters but rather consider many regulator-gene predictions obtained under different hyperparameters. In addition, we subsample the expression data many times to further improve robustness. The final SINGE network is obtained from an ensemble of all of the individual predicted networks using different hyperparameters and cell subsamples (Figure 1).

**Figure 1:**
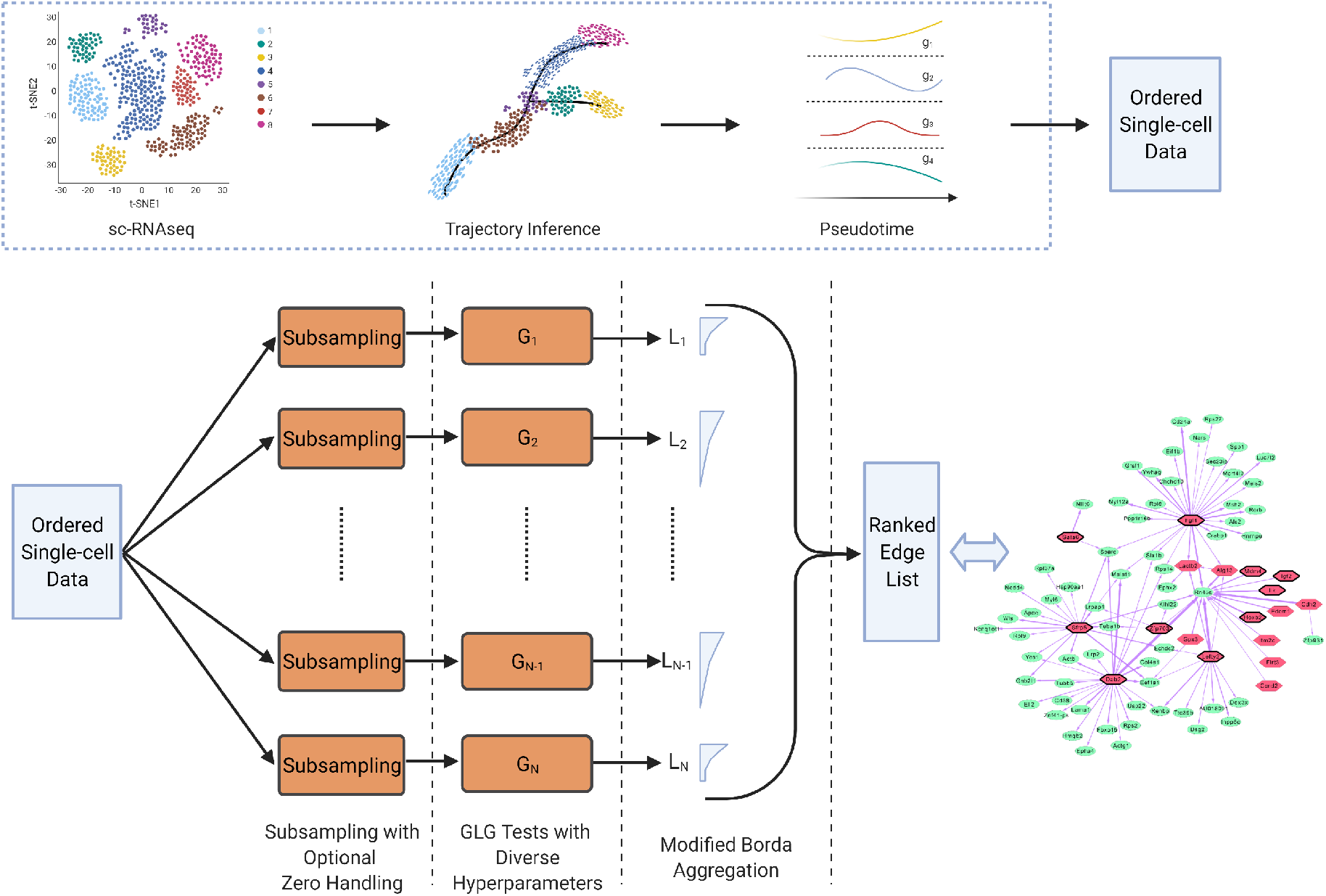
(top) Before running SINGE, trajectory inference methods order single-cell data along the biological process and assign pseudotime values to each cell. SINGE takes the pseudotemporally ordered single-cell gene expression data as input and predicts a ranked list of regulator-target gene interactions. (bottom) SINGE performs multiple GLG tests, each for a different hyperparameter combination and a unique subsample of the ordered data. Optional zero handling removes some of the zero-valued samples for each gene to treat them as missing data. The hyperparameters control sparsity, kernel smoothing, pseudotemporal resolution, and history for the GLG tests. Each GLG test predicts a partial regulatory network, which is converted to a ranked gene interaction list. These preliminary rankings may not include all regulator-target pairs due to the sparsity settings. The individual ranked lists are aggregated into a final ensemble GRN prediction with a modified version of Borda aggregation. Created with BioRender.com.

### 2.2. Network Inference Case Studies

#### 2.2.1. Mouse Embryonic Stem Cell to Endoderm Differentiation

Our first application tracks the differentiation of mouse embryonic stem cells (ESC) to primitive endoderm cells over 72 hours [51]. Matsumoto et al. [15] previously pre-processed this dataset to benchmark their SCODE GRN algorithm. We reuse their processed version of the data, which included expression data for only 100 TFs, 356 cells, and pseudotimes assigned with Monocle. We use this ESC to endoderm differentiation dataset to optimize and tune aspects of the SINGE algorithm, such as the modified Borda aggregation (Section 3.2.5), assessing how well it recovers known regulator-gene interactions that are relevant in mouse embryonic stem cell differentiation from the ESCAPE database [52]. The ESCAPE gold standard is incomplete due to lack of experimental data for many of the relevant TFs (see Section 3.4.1). Therefore, the gold standard only contains an 11 × 99 subset of the 100 × 99 regulator-gene interactions that SINGE scores. SINGE does not score self-edges.

The SINGE regulatory network is a ranked list of scored regulator-gene interactions (Supplementary File 1). SINGE ranks Foxd3, Gli2, and Nanog as the three most influential regulators in the 100-gene subnetwork. To illustrate a GLG-inferred regulatory edge, we consider Pou5f1 as an example target gene. Figure 2 shows that using additional past information from GLG-identified regulators improves the predicted expression trend of Pou5f1 along pseudotime. As more genes are added in decreasing order of the ‘edge weight,’ the predicted expression trends become more accurate.

**Figure 2:**
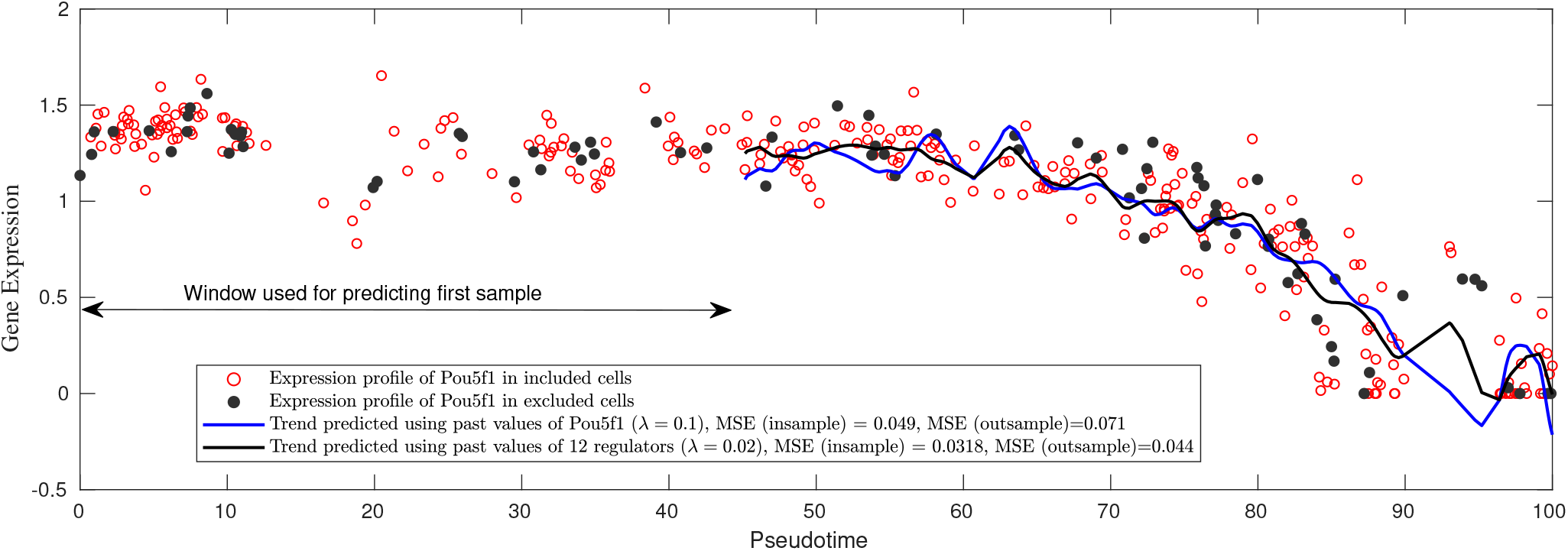
Generalized Lasso Granger example. We show results from individual GLG tests for predicting the regulatory influence on target gene Pou5f1 in the ESC to endoderm differentiation dataset with two values of the sparsity hyperparameter λ. λ = 0.1 selects one regulator, Pou5f1, and λ = 0.02 selects 12 regulators including Pou5f1. We predict expression trends of Pou5f1 using the expression measurements that were included in the GLG model trained using the insample cells. As λ is reduced, we observe a decrease in both the insample mean-squared error (MSE) and outsample MSE of the Pou5f1 expression. Outsample MSE is calculated over the Pou5f1 expression values removed during the subsampling. We also observed a consistent phenomenon for λ = 0.05 (insample MSE = 0.045, outsample MSE = 0.053) and for λ = 0.01 (insample MSE = 0.027, outsample MSE = 0.035), which are not shown in the figure. All reported MSE values are the mean of 10 GLG runs with independent insample/outsample splits.

To assess whether SINGE can match or exceed the state-of-the-art performance after dataset-specific tuning, we compare its predicted GRN with four existing network inference methods: SINCERITIES [27], which uses ridge regression motivated by Granger Causality; SCODE [15], which is based on ordinary differential equations; Jump3 [14], based on decision trees on temporal transcriptomic data; and its predecessor GENIE3 [12], the best performing method in the DREAM4 *In Silico Multifactorial* challenge [53], which does not use temporal information. We emphasize this particular evaluation is not indicative of which method would perform best on new data because of SINGE’s tuning. Nevertheless, SINGE performs similar to the other methods with respect to the average precision (**A**) and much better with average early precision (**E**), which both summarize a precision-recall curve (Figure 3). Average early precision emphasizes the most-confident, top-ranked interactions (Section 3.4.2). Even though SCODE was previously evaluated using this gene expression data [15], it performs worse than random when assessed using the condition-specific ESCAPE gold standard (Section 4.2).

**Figure 3:**
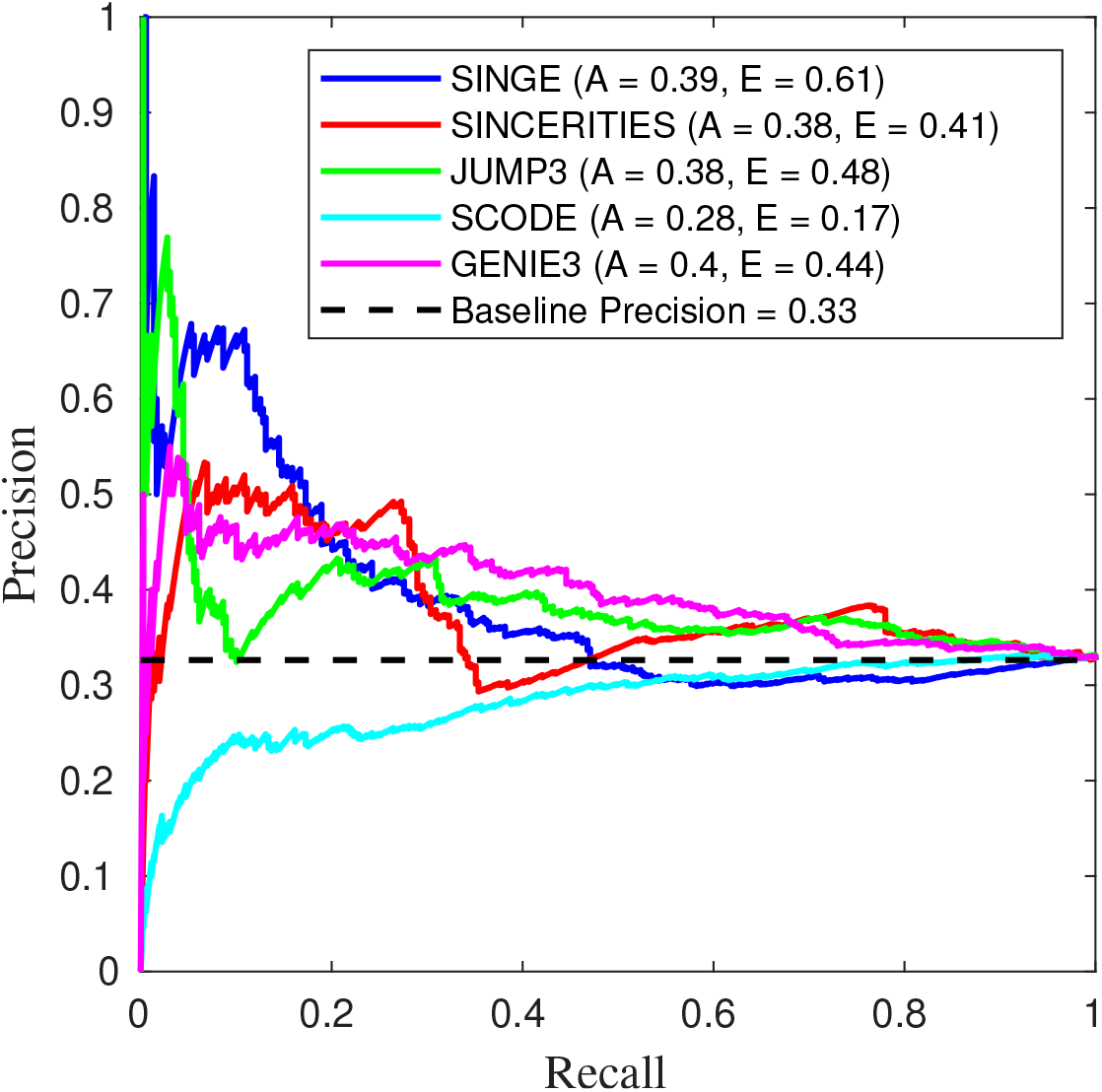
Precision-recall performance of SINGE compared to SINCERITIES, Jump3, SCODE, and GENIE3 when predicting the GRNs involved in primitive endoderm differentiation. The Baseline Prediction is the expected precision obtained by randomly ordering all regulator-gene interactions. SINGE is tied for best place with Jump3 and SINCERITIES in average precision but shows much better average early precision than all other methods. Key: **A** - Average Precision, **E** - Average Early Precision (≤ 0.1 recall).

#### 2.2.2. Mouse Retinoic Acid-driven Differentiation

We further test SINGE on a second dataset that tracks retinoic acid-driven differentiation from mouse embryonic stem cells to extraembryonic endoderm and neuroectoderm cells over 96 hours [54]. SINGE is not tuned for this dataset or the subsequent applications (Sections 2.2.4 and 2.2.5). It uses the same version of the algorithm and hyperparameters from the ESC to endoderm differentiation analysis (Section 2.2.1). We infer a trajectory for the differentiation process using Monocle 2 [25] and select 1886 cells from cell states 1 and 2 (Figure S1 and Section 3.3.2). Monocle 2 also identifies 626 genes whose expression changes substantially as a function of pseudotime, which we use for GRN reconstruction. These genes are not filtered to include only TFs or other known expression regulators. SINGE returns a ranked list of all 626 × 625 possible regulatory relationships, excluding self-edges (Supplementary File 2).

SINGE identifies key regulators reflecting the differentiation trajectory required for mouse embryonic stem cells exiting the pluripotent state, transitioning through the epiblast where lineage segregations take place [54] (Table 1 and Figure 4). We use g:Profiler [55] to identify Gene Ontology (GO) biological process terms that are significantly enriched among the ranked SINGE regulators (Supplementary File 3). This searches for GO terms that are enriched at the top of the ranked list, assessing all possible rank thresholds. The g:Profiler analysis identifies relevant significantly enriched biological processes in the sorted regulator list including cellular response to growth factor stimulus (GO:0071363), cell morphogenesis involved in differentiation (GO:0000904), neuron differentiation (GO:0030182), and additional terms depicted in Table 1.

**Figure 4:**
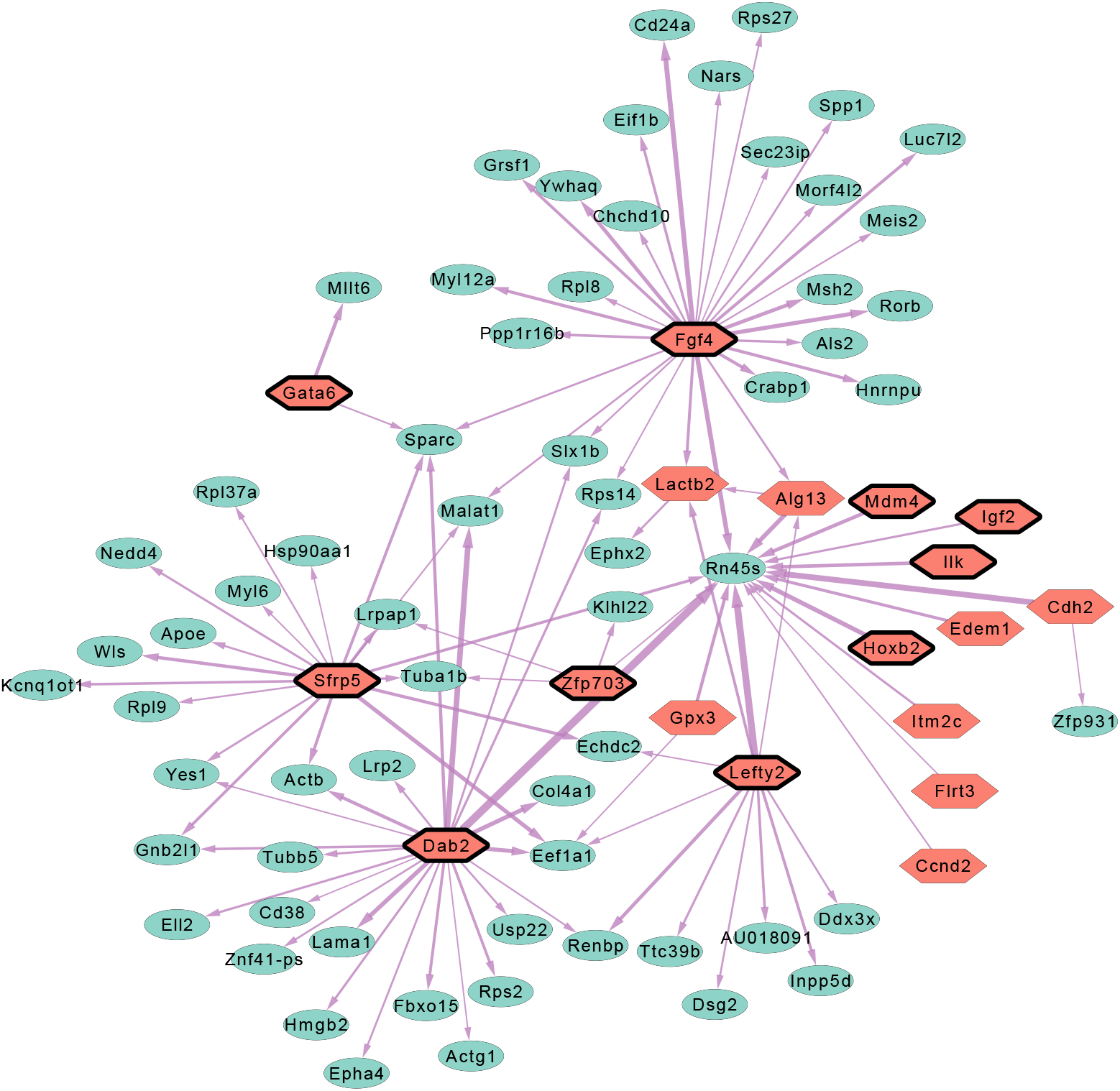
The network obtained from the top 100 edges ranked according to SINGE scores shows 18 unique regulators (hexagonal nodes, the ten with solid boundaries corresponding to known regulators of gene expression listed in Table 1) and 65 unique targets (elliptical nodes). The higher ranked edges are represented by thicker arrows.

**Table 1:**
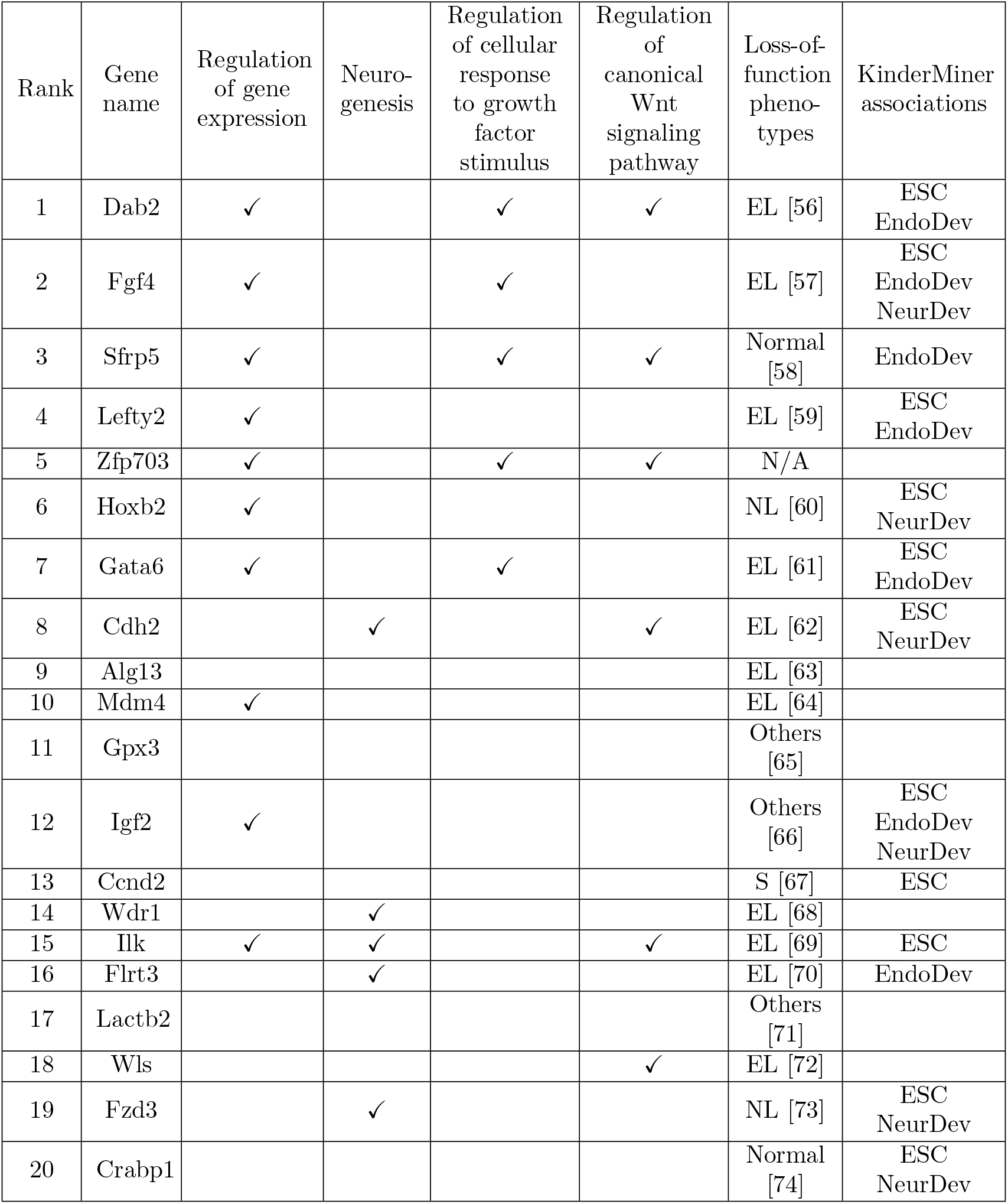
GO biological process terms, loss-of-function phenotypes, and KinderMiner associations related to the top 20 SINGE regulators. **Phenotype key —** EL: Embryonic lethality, NL: Neonatal lethality, S: Sterile, Normal: Homozygous mutant mice are phenotypically normal and fertile, Others: Homozygous mutant mice display other physiological phenotypes, N/A: No knockout mice reported in peer-reviewed studies. **KinderMiner key —** ESC: Embryonic Stem Cells, NeurDev: Neural development, EndoDev: Endoderm development

There are two ways to explore the SINGE predictions in greater detail: the top regulators ranked by SINGE influence (Table 1), which aggregates influence over all target genes, and the top-ranked edges (Figure 4). Table 1 shows the top 20 regulators ranked by SINGE influence. Ten of the top predicted regulators are associated with regulation of gene expression (GO:0010468), as are other regulators with high SINGE influence that are beyond the top 20 (Supplementary File 3). The top 20 regulators also include essential genes that cause embryonic lethality in mouse embryos harboring homozygous null alleles. Others show phenotypes ranging from postnatal lethality to growth retardation (Table 1). Three of the predicted regulators (Alg13, Gpx3, and Lactb2) are known for their roles in metabolic processes but are not known to participate in regulation of early embryonic lineage specification. In addition, KinderMiner [75] text mining reveals significant associations between the top 20 regulators and terms related to this developmental process: ‘embryonic stem cells,’ ‘neural development,’ and ‘endoderm development’ (Supplementary File 4).

Figure 4 illustrates the most-confident 100 regulator-gene edges from the SINGE network, directed from the regulators (hexagons) to the target genes (ellipses). This representative subnetwork comprises 18 unique regulators and 65 unique targets. Fourteen of these regulators are also found among the top 20 regulators by SINGE influence (Table 1), including all 10 known to be associated with regulation of gene expression. The other four regulators participate in one or more high-confidence edges but do not have high aggregate influence. Dab2 and Fgf4 are the most influential regulators overall (Table 1) and hub regulators among the top 100 edges (Figure 4). Fgf4 governs the exit from the pluripotent state. *Fgf4-null* mouse embryonic stem cells resist neural and mesodermal lineage induction [76]. The Fgf/Map kinase signaling pathway plays multiple roles during mouse blastocyst development, and mutations of the signaling components (e.g., Fgf4, Fgfr2, and Grb2) all cause implantation lethality and lack of primitive endoderm development [77]. Moreover, Fgf4 also governs neural induction in embryonic stem cell differentiation at a later stage of development [78]. Rn45s is predicted to be a frequently-regulated target gene. However, these are likely false positive predictions. Rn45s is 45S pre-ribosomal RNA, and its expression levels and variance are much higher than any of the other 625 genes in this dataset.

The predicted GRN in Figure 4 also provides hypotheses for future experimental tests. For example, Meis1 and Meis2 are homeobox proteins that directly regulate Pax6 expression during eye development [79]. SINGE predicts that Fgf4 regulates Meis2. Thus, Fgf4 could potentially act upstream of Meis1 and Meis2 to regulate Pax6 expression, contributing to neuroectoderm differentiation [80]. Other key primitive endoderm regulators are also highlighted in SINGE predictions, such as Gata6, a transcription factor necessary and suf-ficient for primitive endoderm lineage differentiation and establishment of extraembryonic endoderm cell lines [81]. Dab2, Sfrp5, Lefty2, and Igf2 are all expressed in the primitive endodermal lineages, including visceral endoderm and extraembryonic endoderm cell lines [82–86].

Inspecting the regulator and target expression trends can build confidence in the predicted interactions. For example, in the predictions Dab2→Yes1 and Fgf4→Meis2, we observe that the target gene expression is negatively correlated with the past values of the regulator’s expression (Figure S2), which may indicate these predictions are worth further investigation. In other cases like Dab2→Rn45s, there is no obvious relationship between the regulator and target expression. As noted above, this is likely a false positive prediction due to Rn45s’s outlier expression levels and role as a pre-ribosomal gene.

Many expected GO terms and regulators are represented in Table 1 and Figure 4. However, classic neuroectoderm regulators like Sox1, Nes, and Pax6 [54] are missing because they are excluded from the limited shortlist of genes in the SINGE input. We only run SINGE on the top 626 significantly differentially expressed genes along the differentiation trajectory detected by Monocle 2.

#### 2.2.3. Retinoic Acid-driven Differentiation ESCAPE Evaluation

The retinoic acid-driven differentiation study can be used to benchmark the relative performance of SINGE and other network inference methods because none of the methods, including SINGE, were optimized or tuned based on the ESCAPE evaluation results. Figure 5 shows the precision-recall performance of SINGE, SINCERITIES, Jump3, SCODE, and GENIE3 when ranking edges in the 626-gene network. Due to Jump3’s runtime, we run it on a reduced dataset (Section 3.4.3), which may impact its performance. As with the ESC to endoderm differentiation dataset, the ESCAPE database had only partial information (12 regulators), thus limiting the gold standard to a submatrix of 12 × 625 possible edges. SINGE is the best method overall in terms of average precision and average early precision (tied with Jump3) (Figure 5). Jump3 is effectively tied with SINGE for average early precision but has near-random precision for recall > 0.2. SINCERITIES prioritizes ESCAPE gold standard interactions well at the top of its ranked list, but the performance degrades quickly. GENIE3 and SCODE are worse than random. The performance depends on the type of regulator-gene interaction in the ESCAPE database. SINGE can recover loss-of-function or gain-of-function (*lof/gof*) relationships but struggles to identify ChIP-based protein-DNA binding interactions (Figure S3). In contrast, Jump3 and GENIE3 recover ChIP-based protein-DNA binding interactions quite well but struggle to identify *lof/gof* relationships.

**Figure 5:**
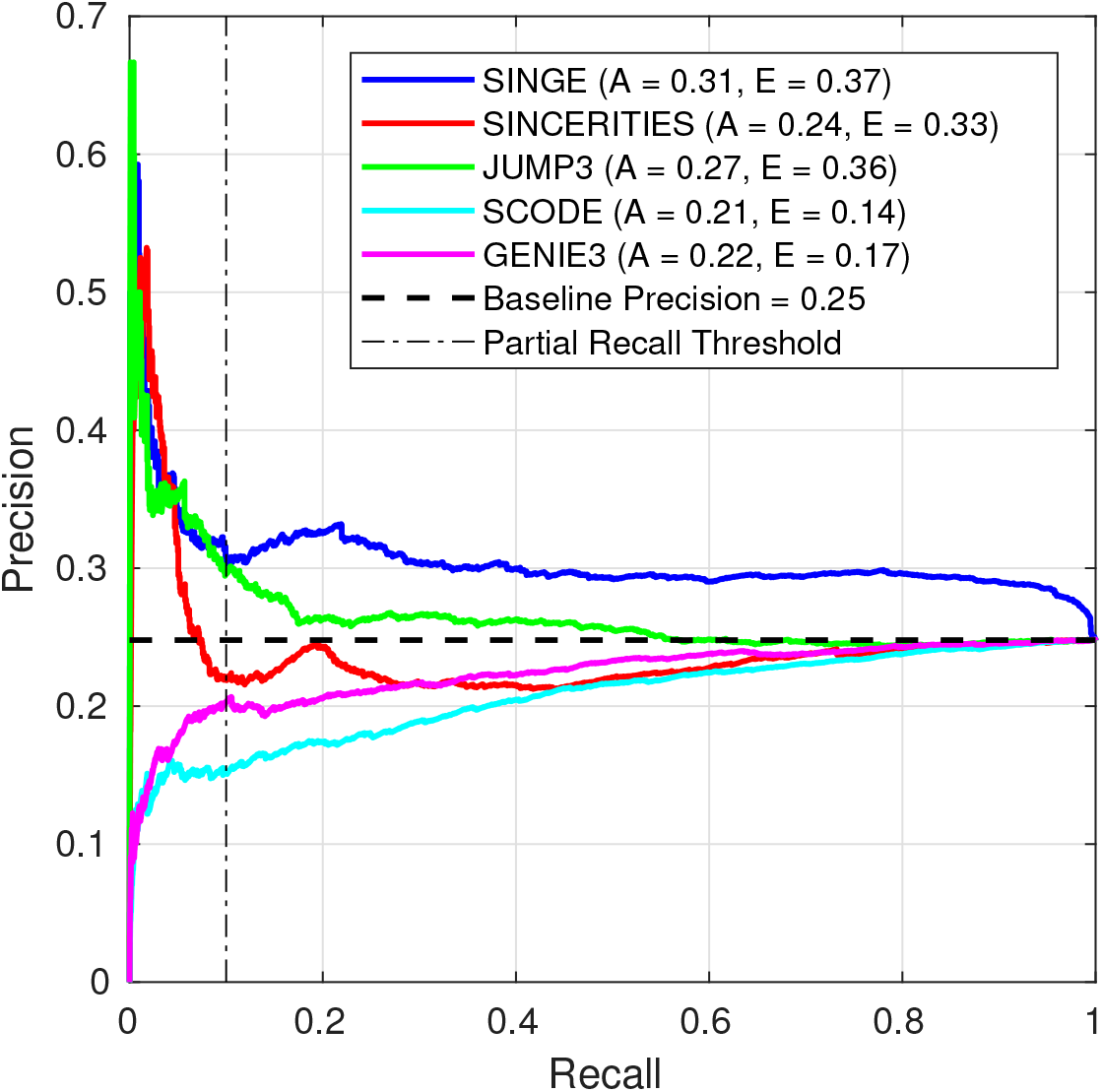
Precision-recall performance of SINGE, SINCERITIES, Jump3 (which uses a reduced data set), SCODE, and GENIE3 when predicting a 626-gene retinoic acid-driven differentiation regulatory network [54]. SINGE is tied for best with Jump3 in average early precision but is better in average precision. Key: **A** - Average Precision, **E** - Average Early Precision (≤ 0.1 recall).

Visualizing the expression trends over pseudotime can illustrate the types of errors SINGE makes with respect to the ESCAPE gold standard. For example, the interaction Esrrb→Actb was detected with ChIP but is not part of the ESCAPE’s *lof/gof* dataset. There is no apparent lag between the expression trends of the regulator and target (Figure S4). This edge was ranked highly by SCODE but not by SINGE, which searches for lagged expression dependencies by design.

A regulator-specific evaluation partially explains the overall precision-recall performance of the GRN methods and demonstrates that it can be somewhat misleading. Figure 6 shows the average precision and average early precision with respect to each regulator in the ESCAPE database. These metrics are obtained from the regulator-specific precision-recall curves in a similar manner to the average precision and average early precision obtained in Figure 5. Recall that the average early precision is the average precision in the early part (recall ≤ 0.1) of the precision-recall curve. The regulator-specific average precision of all five methods is at or below random for most regulators, with a few exceptions.

**Figure 6:**
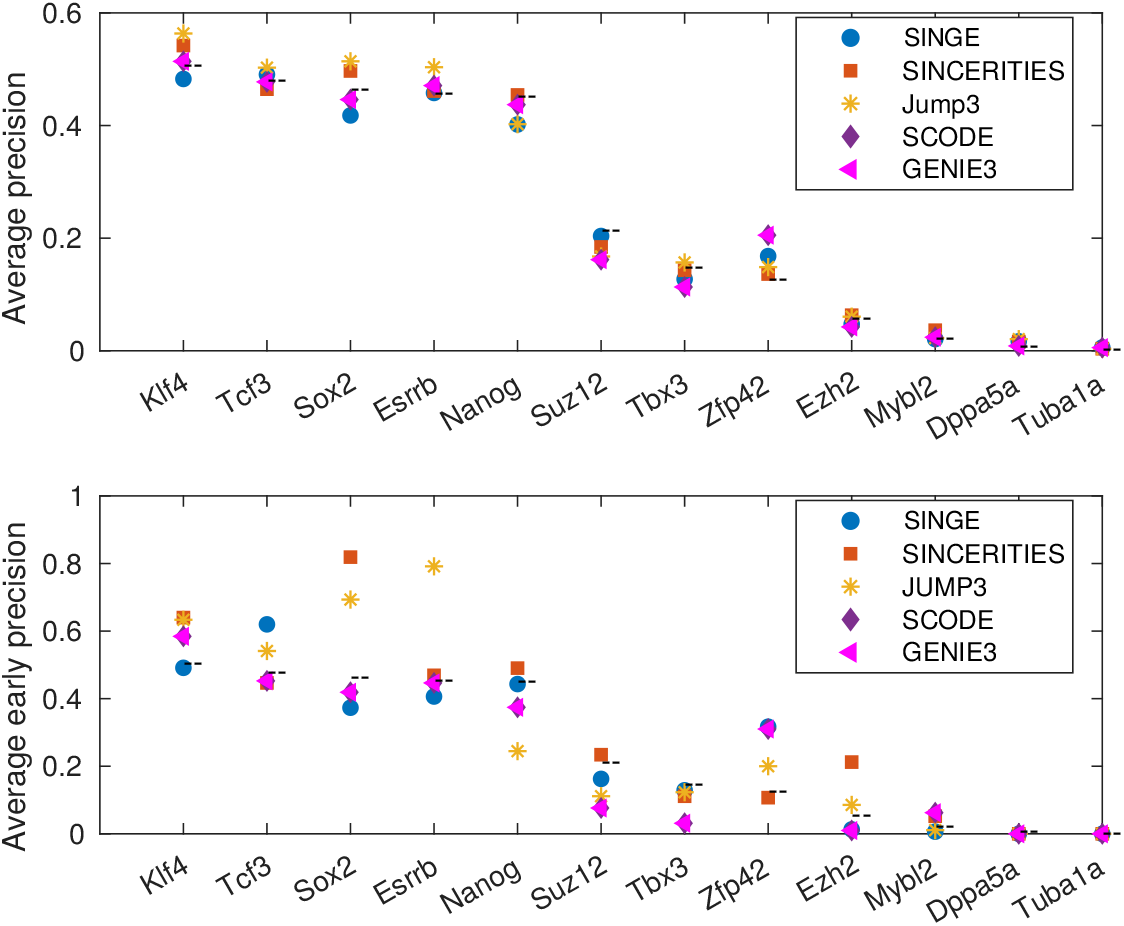
Average precision and average early precision evaluated for individual regulators in the ESCAPE database. The dashed line (– –) indicates the expected performance of a random ranking, given by the ratio (total number of true outgoing edges)/(total number of genes - 1). Regulator-specific performance of all five methods is at or below random for most regulators.

Because some regulators are more prevalent in the ESCAPE gold standard than others, the overall precision-recall curve is influenced by both the regulator-specific precision and the relative ordering of the regulators in the ranked edge list. We can sort these 12 regulators in decreasing order by their number of outgoing edges in the ESCAPE gold standard, which is a proxy for the regulator’s influence on the evaluation, and generate boxplots of the regulator-specific edge ranks in the GRNs (Figure 7). SINGE ranks outgoing edges from ESCAPE’s most prevalent regulators (Klf4 and especially Tcf3) higher on average than the regulators with fewer target genes (Dppa5a and Tuba1a). The distributions of rankings from SINCERITIES and Jump3 are widely dispersed for each regulator. Meanwhile, SCODE and GENIE3 rank edges from the regulators with fewer outgoing edges higher than those with many target genes, contributing to their poor overall performances.

**Figure 7:**
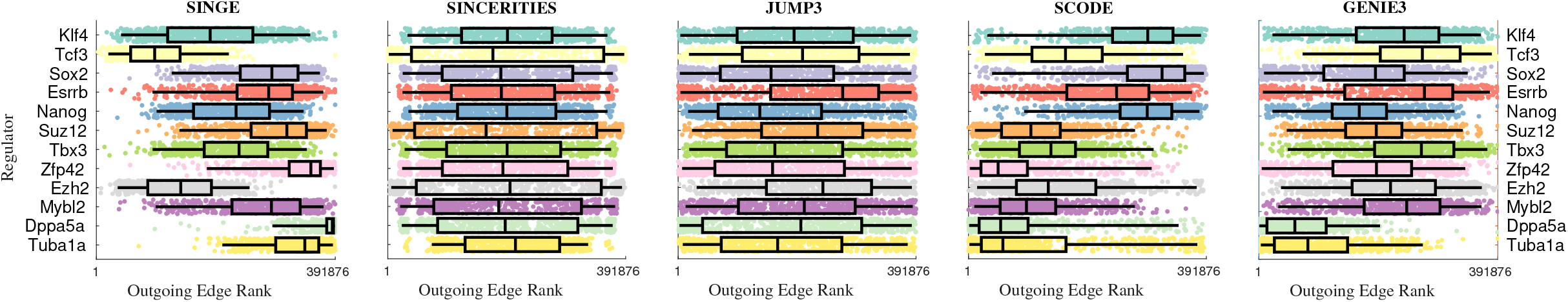
The ranking of regulator-specific interactions has a strong effect on the overall precision-recall curve (Figure 5). The boxplots show the outgoing edge ranks for each regulator in each predicted GRN, in decreasing order of regulator prevalence in the ESCAPE database. Ranking regulator-gene interactions involving the predominant ESCAPE regulators (e.g., Klf4) above those involving the less frequent ESCAPE regulators (e.g., Tuba1a) improves the precision-recall performance, and the converse is also true.

These regulator-specific results provide insights into Figure 5. SINGE’s relatively high average early precision is influenced by how it ranks regulators in accordance with their prevalence in the ESCAPE database. On the other hand, Jump3 ranks all regulators uniformly but has better than random average precision on multiple individual regulators, such as Sox2 and Esrrb. When evaluated from the perspective of the individual regulator-specific precision-recall performance, all five methods perform at near random for most regulators (Figure 6), but this does not necessarily translate into near-random precision-recall performance for the entire GRN (Figure 5). This is because some of the methods rank certain regulator-specific interactions above others (Figure 7) either to their benefit (SINGE) or to their detriment (SCODE and GENIE3).

#### 2.2.4. Mouse Bone Marrow Mesenchyme to Erythrocyte Differentiation

As an additional SINGE case study, we choose an scRNA-seq dataset from the Mouse Cell Atlas profiling the heterogeneity of adult mouse bone marrow [87]. This dataset helps assess SINGE’s scalability to more genes (3025 genes) and cells (3105 cells). We also generate the pseudotimes from an alternative trajectory inference algorithm, Embeddr [88]. We hypothesize that the bone marrow scRNA-seq data should shed light on regulators for hematopoiesis or its associated diseases. Indeed, SINGE identifies relevant regulators in this context. For example, Asxl2 and Rtel1 are among the top 20 regulators. Asxl2 knockout mice displayed a phenotype that skewed the differentiation potential of hematopoietic stem cells and cause myeloid-lineage cancer [89]. In humans, Asxl2 mutation is known to associate with acute myeloid leukemia [90]. Rtel1 is a DNA helicase important for protecting telomeres. Its mutations are associated with a number of clinical phenotypes related to bone marrow failure [91, 92]. Although other top regulators do not have direct experimental data to show their role involved in hematopoiesis, the SINGE GRN (Supplementary File 5) provides a base for future investigations.

#### 2.2.5. Dyngen Simulation Evaluation

We use `dyngen` [93] to evaluate GRN inference on simulated single-cell datasets. We simulate single-cell gene expression data from a biological process with a linear trajectory. This simulated dataset has 1000 cells generated from a regulatory network with 140 genes — 25 TFs, 15 housekeeping genes, and 100 target genes. We use SINGE and the four other GRN algorithms to infer networks from this dataset. We first evaluate the precision-recall performance of each inferred network using the known direct regulatory interactions (precisionrecall curves available at https://github.com/gitter-lab/SINGE-supplemental). Most methods, including SINGE (Average precision = 0.0082, Average early precision = 0.0048), perform as poorly as the random baseline precision of 0.0084, with Jump3 performing best (Average precision = 0.015, Average early precision = 0.032). If we include indirect regulator-gene interactions in the gold standard, the precision-recall performance of SINGE improves, surpassing the random baseline and other GRN methods, which remain near or worse than random. Thus, in the `dyngen` simulation, SINGE performs poorly at distinguishing direct gene interactions from indirect ones. This is in part because the simulated gene expression trends of a regulator’s direct and indirect targets can be quite similar, as exemplified by the cascade from B1_TF1 to Target1 to Target48 (https://github.com/gitter-lab/SINGE-supplemental).

### 2.3. Analyzing Features of the SINGE Workflow

We use the retinoic acid-driven differentiation dataset to perform in-depth analyses of various SINGE features. This is the largest available real dataset from our case studies that has a gold standard available.

#### 2.3.1. Effects of Subsampling and Zero Handling

SINGE’s ensembling can improve performance by supporting subsampling and zero handling. Because the core GLG test is compatible with irregular time series, we can create randomly subsampled time series from each gene’s expression data to generate multiple instances of the original dataset. In these experiments, subsampled replicates are created by removing individual expression data samples with probability of removal 0.2. The default SINGE setting uses 10 subsampled replicates per hyperparameter combination. Figure 8 shows the effects of increasing or decreasing the number of replicates. For both average precision and average early precision, there are only modest performance improvements when running SINGE with more than 10 replicates. The average precision decreases when fewer replicates are used, but the changes are not substantial. On the other hand, running SINGE with only two replicates improves the runtime considerably but leads to a more notable decrease in average early precision. Reducing the number of replicates from 10 to five maintains similar average early precision and still reduces the runtime.

**Figure 8:**
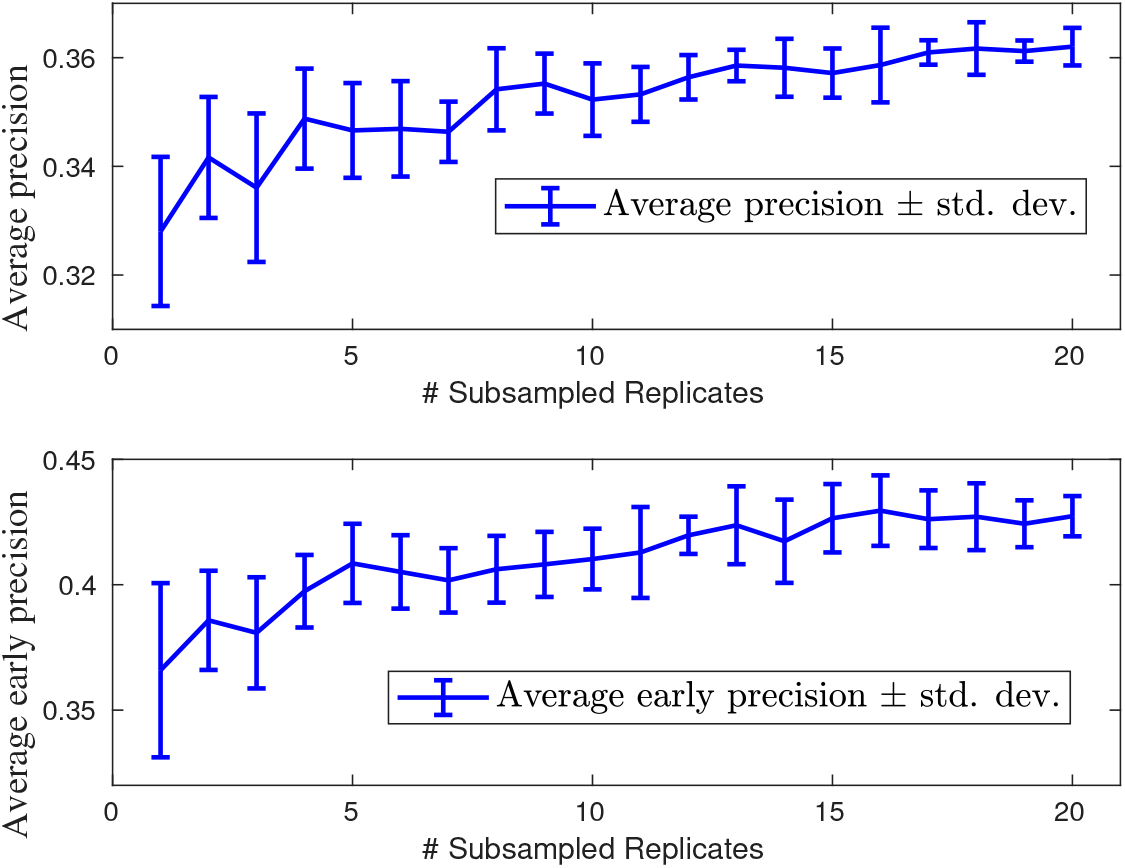
Effect of number of subsampled replicates for each hyperparameter combination in the SINGE ensemble on the retinoic acid-driven differentiation dataset. Each subsampled replicate is generated by arbitrarily dropping samples corresponding to individual genes in each cell with probability 0.2.

The support for irregular time series also allows us to remove zero-valued data points corresponding to technical dropouts. The true dropout probability is gene dependent and can be estimated by methods like SCONE [94]. As a proof of concept of SINGE’s support for zero handling, we incorporate a simpler strategy that uses a constant dropout probability hyperparameter *prob-zero-removal* for all genes. For each GLG instance, we remove zero-valued expression samples (and their corresponding timestamp) from each gene’s expression series with a user-specified constant probability for each zero value.

Figure 9 shows SINGE’s precision-recall summaries as the value of *prob-zero-removal* increases. As more zeros are dropped from the dataset, the average precision and especially the average early precision are only marginally affected. Because dropping samples reduces the size of the regression problem, zero-dropping potentially could be used to effectively reduce the size of the regression problem for large but extremely sparse datasets without negatively impacting GRN inference. Filtering too many zeros in such an arbitrary manner could remove genuine zero expression values along with the dropouts. We currently recommend using SINGE without dropping zeros unless it is required to speed up analysis of large datasets. We will explore directly supporting gene-dependent dropout for improving the precision-recall performance (Section 4.3).

**Figure 9:**
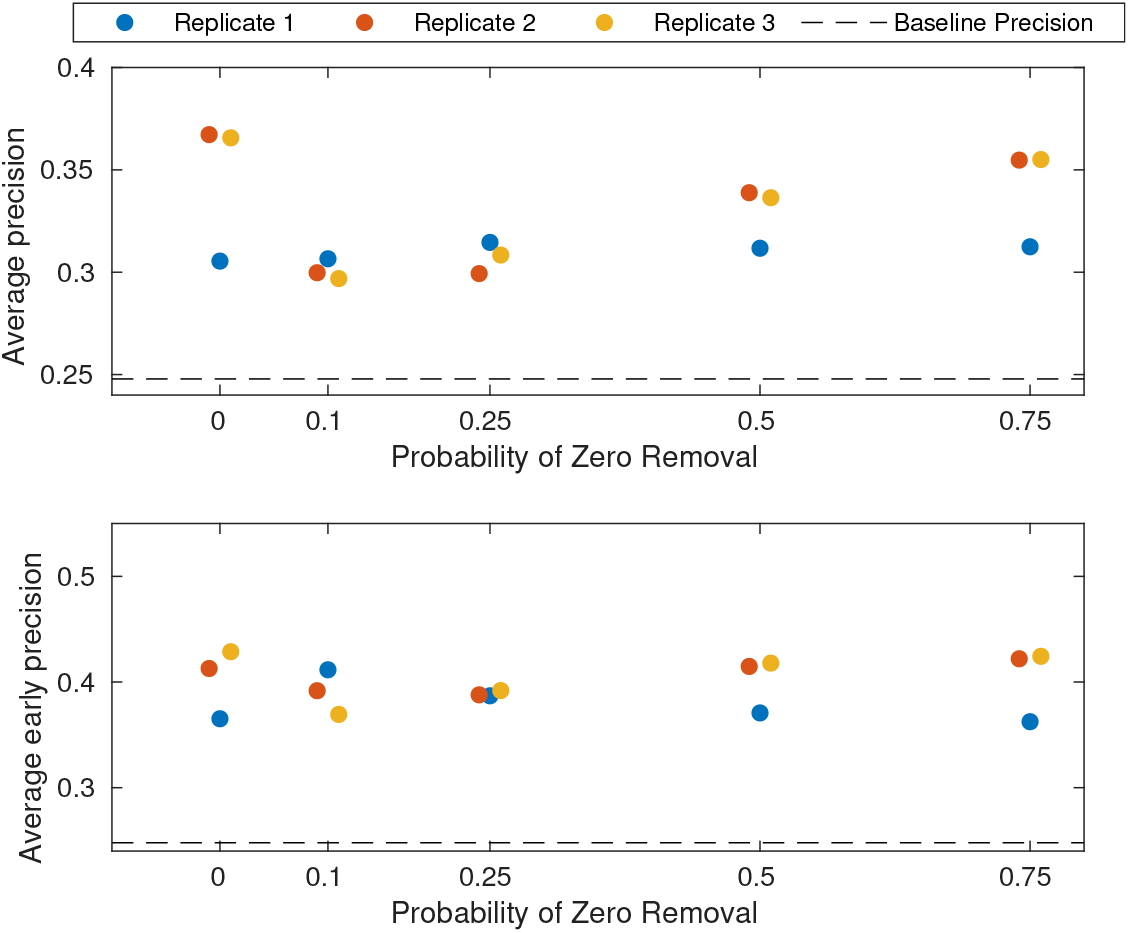
Effect of zero removal on SINGE performance for multiple values of *prob-zero-removal,* the probability of removing a zero value. SINGE ensembles the results from 10 zero-filtered replicates on the retinoic acid-driven differentiation dataset.

#### 2.3.2. Benefits of Ensembling

The optimal GLG parameters that best identify causal relationships between two genes can vary from gene to gene and for different biological processes. In the absence of prior information about the regulatory network, it is difficult to set optimal hyperparameters for the GLG test. Furthermore, it is also plausible that different transcriptional regulators have different kinetics and consequently different optimal hyperparameters.

SINGE attempts to overcome this with an ensemble of hyperparameters, aggregating the results to obtain the final SINGE score of each GRN edge (Section 3.2.5). Figure S5 compares the performance of individual GLG hyperparameter combinations to the complete ensembled SINGE GRN for the retinoic acid-driven differentiation dataset. Although the ensembled SINGE network does not have the best average precision or average early precision, it performs better than the majority of the individual hyperparameters. Ensembling reduces the risk of choosing a single set of hyperparameters that would perform poorly for a particular dataset. Inspecting the performance of the GRNs for individual hyperparameters (Figures S6 and S7) shows that the sparsity hyperparameter λ has the strongest impact.

#### 2.3.3. Assessing whether Pseudotimes Improve GRN Reconstruction

We assess the impact of using assigned cell order and pseudotime values on the performance of the three methods designed to reconstruct GRNs from pseudotemporal single-cell gene expression — SINGE, SINCERITIES, and SCODE. We exclude GENIE3 and Jump3 because the former does not use any sort of cell ordering information, and the latter uses only the cell ordering but not pseudotime values. For this assessment, we create variants of both the ESC to endoderm differentiation and retinoic acid-driven differentiation datasets as described below:

- *Pseudotime:* The default mode using ordered cells with Monocle or Monocle 2 assigned pseudotimes.
- *Order Only:* Obtained from the *Pseudotime* dataset by removing the assigned pseudotime values but maintaining the cell order. The cells are assumed to be regularly-spaced along the trajectory.
- *Rand. Order (3):* Three replicates obtained from random permutation of the regularly-spaced cells from the *Order Only* variant. The randomized data have neither pseudotime annotations nor ordering information from the original dataset.

If estimated pseudotimes contribute high-quality information for GRN reconstruction, the three GRN methods should have highest performance on the *Pseudotime* dataset, with less accurate predictions from the *Order Only* and *Rand. Order* datasets.

Figure 10 shows the average precision and average early precision of SINGE, SINCERITIES, and SCODE when run on the three variants of each dataset above. For variants of the ESC to endoderm differentiation dataset, only SINGE’s performance decreases substantially for the *Rand. Order* dataset as expected. Its performance on the *Order Only* dataset is only slightly worse than the original *Pseudotime* dataset. SINCERITIES is less consistent on the *Rand. Order* datasets, with some randomized cell orders providing better GRNs than the real *Order Only* or *Pseudotime* datasets. SCODE performs poorly even on the original *Pseudotime* dataset (Figure 3) so we cannot draw strong conclusions from its performance trend across the dataset variants.

**Figure 10:**
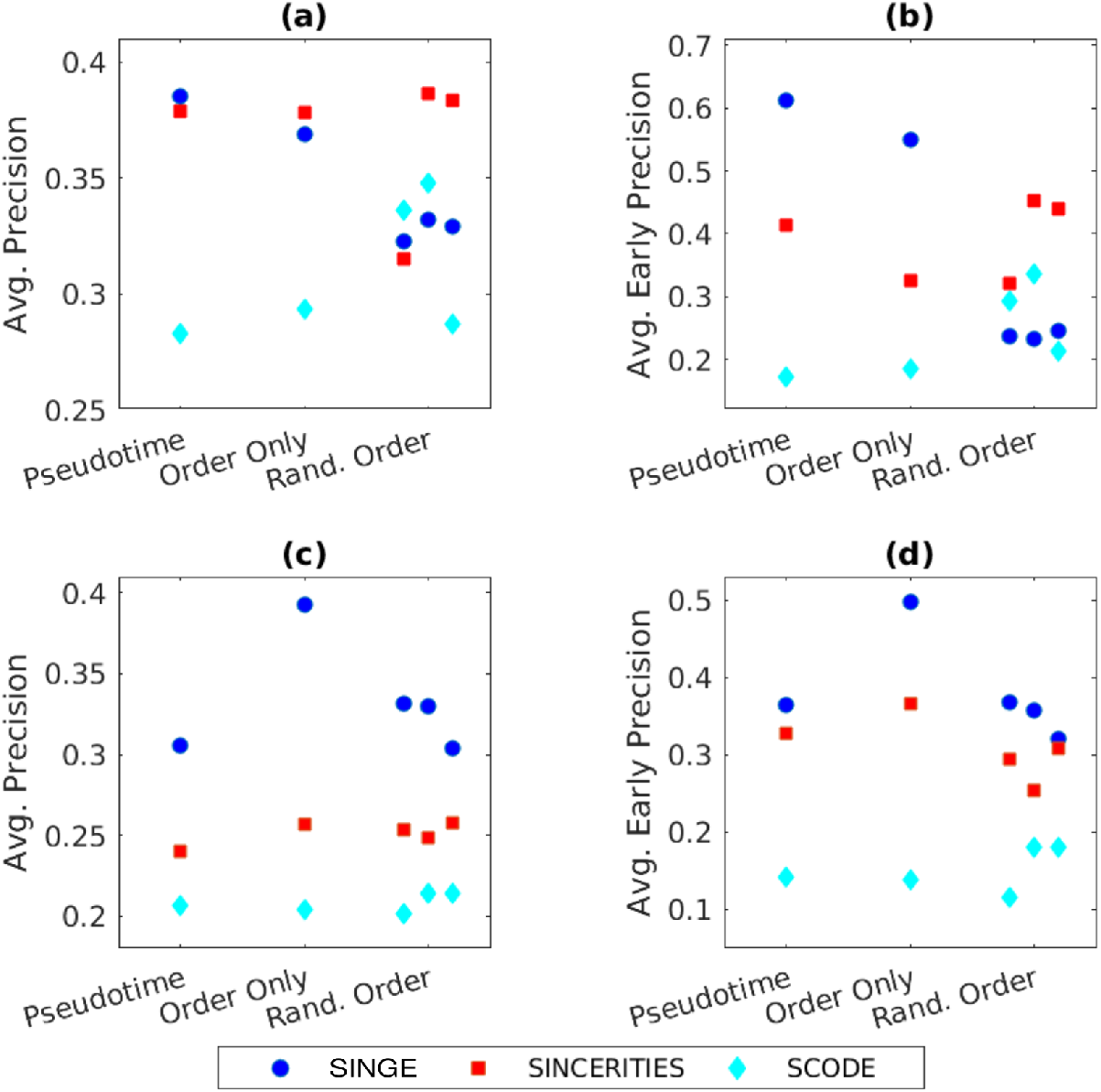
Effects of pseudotimes and cell ordering on the performance of SINGE, SINCERITIES, and SCODE. (a) and (b) show the performance of the three methods when analyzing variants of the ESC to endoderm differentiation dataset. Average precision metrics using Monocle pseudotimes are comparable to those using *Order Only.* (c) and (d) show the performance of the three methods on variants of the retinoic acid-driven differentiation dataset. SINGE's average precision metrics degrade substantially when using Monocle 2 pseudotime values when compared to *Order Only.*

On the other hand, for variants of the retinoic acid-driven differentiation dataset SINGE still outperforms SINCERITIES and SCODE in all cases, but the performance trend does not follow the expected pattern. SINGE shows higher performance on the *Order Only* dataset in which the pseudotime values are removed. Its performance for the *Rand. Order* variants is worse than the *Order Only* dataset but comparable to the *Pseudotime* dataset. SINCERITIES has similar performance on the *Pseudotime* and *Order Only* datasets, with a slight improvement for *Order Only*. SCODE again performs poorly in all cases. Because the performance improves for both SINGE and to a lesser extent SINCERITIES when only the cell ordering is used, one possible explanation is that the Monocle 2 pseudotime values are low fidelity, counteracting any potential benefits from the additional information. Indeed, further analysis of the regulator-specific performance of these three methods using the *Order Only* dataset (Figure S8) shows that the regulator-specific average precision and average early precision metrics of SINGE and SINCERITIES improve compared to the *Pseudotime* dataset (Figure 6). Like Jump3, which does not use the pseudotime values, these two methods now have substantially better than random average early precision for several regulators. Similarly, the SINGE and SCODE average rankings of the outgoing interactions from the regulators using the *Order Only* dataset better match the regulators’ prevalence in the ESCAPE database (Figure S9). Regulators with more interactions in ESCAPE tend to have higher rankings in these predicted GRNs. These two phenomena combine to improve SINGE’s overall precision-recall curve with respect to its *Pseudotime* dataset performance and those of other GRN methods for either form of the dataset (Figure S10).

We further investigate the relationship between the quality of the SINGE-inferred GRN and the trajectory inference method used to generated pseudotimes. We infer another trajectory from the retinoic acid-driven differentiation dataset using PAGA Tree [95] in the `dynverse` environment (Section 3.3.2). We limit our study to the longest branch of the trajectory, which has 2631 cells, of which 737 cells are common with the Monocle 2 branch (Figure S11). Thus, the cell populations in the two test cases have some overlap yet contain cells exclusive to each dataset.

Figure 11 evaluates the precision-recall performance of the network inferred using SINGE version 0.3.0 and the PAGA Tree trajectory with *Pseudotimes* and *Order Only*. For reference, we add the original SINGE (version 0.1.0) precision-recall curve using the Monocle 2 inferred trajectory from Figure 5 as well as the precision-recall curve for the Monocle 2 *Order Only* dataset from Figure S10. To capture any effects due to the software changes between versions 0.1.0 and 0.3.0, we rerun SINGE 0.3.0 on the Monocle 2 trajectory.

**Figure 11:**
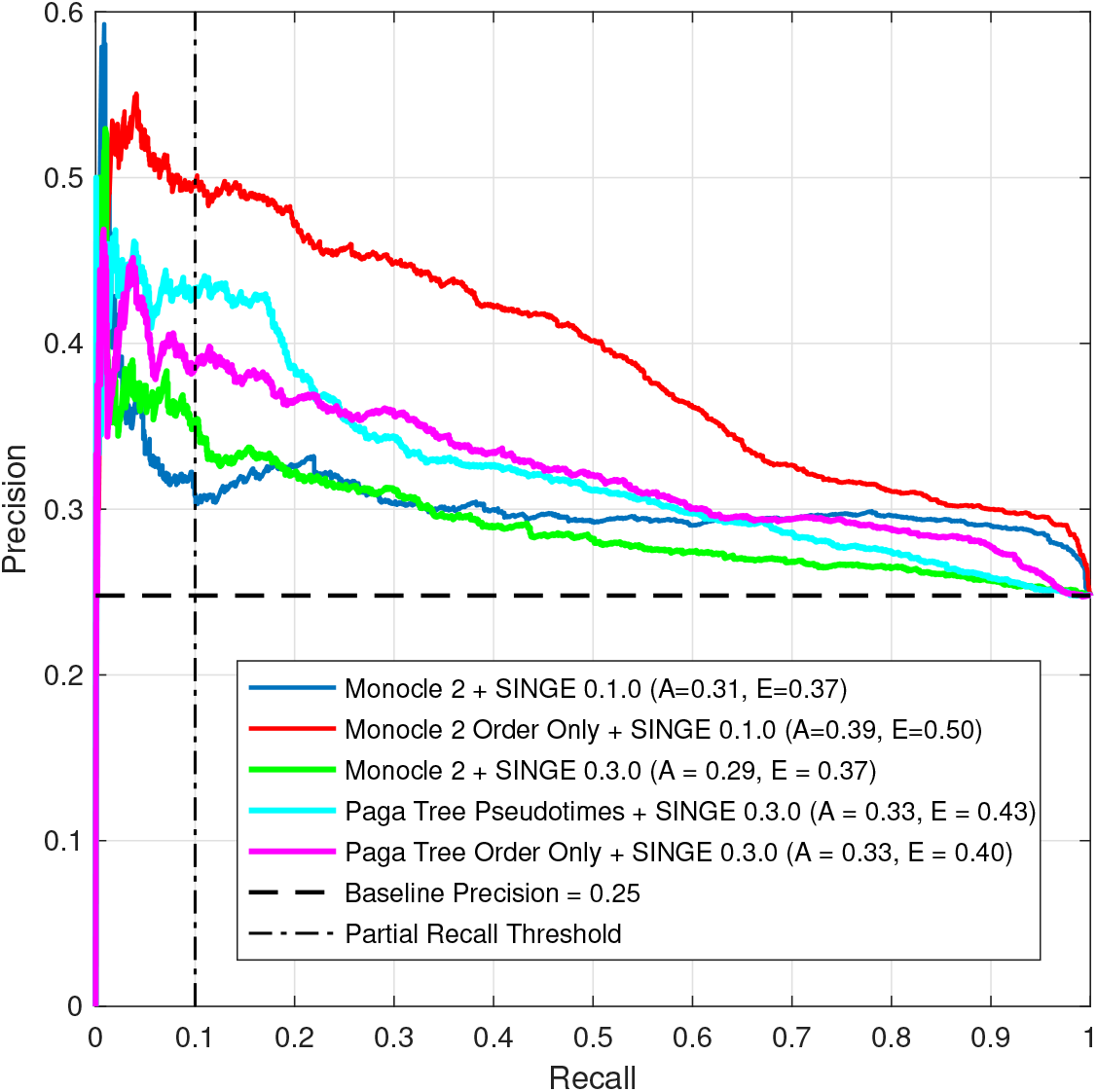
Precision-recall comparison of SINGE using *Order Only* and *Pseudotimes* datasets from Monocle 2 and PAGA Tree trajectories. The comparison of SINGE versions 0.1.0 and 0.3.0 on the Monocle 2 *Pseudotimes* dataset is used to confirm performance consistency across software updates.

The quality of the inferred GRN depends upon the type and quality of the pseudotimes. Importantly, the networks inferred with the original *Pseudotimes* are not always better than the *Order Only* version. The SINGE GRN obtained with Monocle 2 *Order Only* is substantially better than all others. For PAGA Tree, the performance differences between *Pseudotimes* and *Order Only* are negligible. The two SINGE results on Monocle 2 *Pseudotimes* are also comparable. The minor differences in the precision-recall curves for SINGE versions 0.1.0 and 0.3.0 can be attributed to randomization in the algorithm and efficiency-improving optimizations.

### 2.4. Computational Runtime

We designed SINGE to take advantage of the high-throughput computing resources that are readily available to computational researchers, such as the Open Science Grid [96]. We compare the GRN methods’ runtimes on the retinoic acid-driven differentiation dataset. SCODE and SINCERITIES require the least computational resources. It was possible to run them on a single workstation with a 64-bit Intel i5-4590 CPU and 8 GB RAM. Specifically, on this workstation, the SCODE algorithm with 100 repetitions requires approximately 6 hours to complete, whereas the SINCERITIES algorithm takes approximately 111 hours.

In contrast, both SINGE and Jump3 require more varied and extensive computing resources. In the case of Jump3, inferring the GRN from 626 regulators to one target gene takes between 11 minutes to 74 hours to run, with an average runtime of 21.7 hours. This is repeated for each target gene. In a typical application, SINGE uses 100 different hyperparameter settings on 10 subsampled expression datasets. Running SINGE for five λ hyperparameter values on one subsampled replicate takes 6.09 hours (Table 2). The entire SINGE workflow for all hyperparameters and replicates requires 1219.4 hours. However, both Jump3 and SINGE are highly parallelizable. We deployed them on our local high-throughput computing cluster using HTCondor [97], which connects to the Open Science Grid [96]. In this high-throughput setting we can run the entire SINGE algorithm in 36 hours and the Jump3 algorithm in 72 hours.

**Table 2:** Computational runtime of SINGE for datasets of various sizes. SINGE versions 0.3.0 and 0.5.0 have performance improvements not present in version 0.1.0. ‘Average runtime for all λ values’ is calculated as the average compute time taken to run the GLG test for all λ values. SINGE version 0.1.0 runs independent GLG tests for each λ, but starting with version 0.3.0 it obtains the GLG test results for all λ values in one batch by using glmnet's warm-start functionality. ‘Parallel runtime' is calculated using the longest individual runtime for a GLG test. This is the approximate SINGE runtime in the hypothetical scenario where all GLG tests are run in parallel simultaneously. ‘Serial runtime' is calculated as the aggregate compute time of all GLG tests. This is the approximate SINGE runtime in the hypothetical scenario where it is run serially on a single machine corresponding to the average node in the high-throughput computing pool.

| Dataset | Replicates | Genes | Cells | SINGE version | Average runtime for all $\lambda$ values (hours) | Parallel runtime (hours) | Serial run-time (hours) |
| --- | --- | --- | --- | --- | --- | --- | --- |
| Retinoic Acid-driven Differentiation with Monocle | 10 (default) | 626 | 1886 | 0.1.0<br>0.3.0 | 19<br>6.09 | — | 3813<br>1219.4 |
| dyngen dataset | 10 | 140 | 1000 | 0.3.0 | 0.083 | 0.36 | 16.77 |
| Large dyngen dataset | 10 | 140 | 20000 | 0.3.0 | 9.52 | 32.07 | 19038 |
| Retinoic Acid-driven Differentiation with PAGA Tree | 10 | 626 | 2540 | 0.3.0 | 7.83 | 22.36 | 1566.3 |
| Bone marrow dataset (all genes as regulators) | 10 | 3025 | 3105 | 0.3.0 | 23.79 | 62.68 | 4759.2 |
| Bone marrow dataset fast mode (149 TFs as regulators) | 2 | 3025 | 3105 | 0.5.0 | 5.19 | 9.12 | 207.8 |

SINGE can also be configured to run on a single workstation with appropriate changes to the hyperparameters. For example, a complete SINGE run on the bone marrow dataset takes 18 hours 42 minutes using a dedicated 16-core server with Intel Xeon Silver 4110 processors. For this run, we limit the number of regulator genes to only 149 transcription factors (identified using the AnimalTFDB 3.0 database [98]) and reduce the subsampled replicates per hyperparameter combination from 10 to two to reduce the runtime.

## 3. Methods

SINGE infers the GRN that underlies a biological process by aggregating ranked edge lists obtained from an ensemble of Generalized Lasso Granger tests conducted on ordered single-cell transcriptomic data (Figure 1). The GLG test is a kernel-based generalization [47] of the Lasso Granger Causality test to facilitate the analysis of causal relationships between irregular time series obtained from a linear stationary vector autoregressive (VAR) model. We first describe the GLG test and then the complete SINGE algorithm.

### 3.1. Generalized Lasso Granger Test

The GLG test is used to discover temporal causal networks from irregularly-spaced time series data based on concepts of Granger Causality. In GRN inference, the time series correspond to temporal gene expression measurements. Assume *P* regularly-spaced time series *x*_1_, *x*_2_, …, *x_p_* are obtained at timestamps {*t*} ≐ 1,2,…, *T*. These time series are assumed to be governed by a linear and stationary VAR process such that

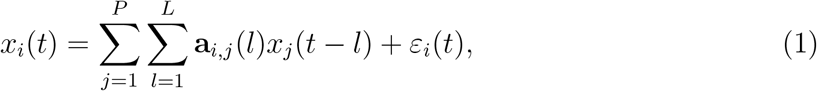

for *i* = 1, 2,…, P, where **a**_*i,j*_(*l*) corresponds to the *l*-th lagged coefficient from source time series *x_j_* to target time series *x_i_* and *ϵ_i_*(*t*) is measurement error, represented by independently distributed Gaussian random variables. More generally speaking, the unknown *P × P* × *L* matrix a comprising L lagged coefficient matrices a(1), a(2),…, a(*L*) represents the evolutionary mechanism of *x*_1_, *x*_2_,…, *x_p_*.

The Lasso Granger Causality test [48] for an individual regularly-spaced target series x¿ is characterized by the optimization problem

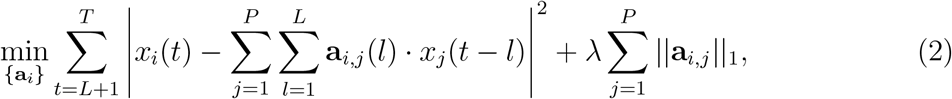

and provides a sparse estimation of the *P × L* coefficient matrix a_*i*_ representing the VAR process that relates each source series *x_j≠i_* to the target series *x_i_*. Specifically, if elements of the *j*-th column a_*i,j*_ of the matrix are statistically significant, then we claim that *x_j_* Granger-causes *x_i_* (represented by *j → i*). The Lagrange multiplier λ dictates the sparsity of the learned matrix a_*i*_.

The GLG test proposed by Bahadori and Liu [47] is a kernel-based modification of Equation 2 to facilitate the analysis of irregular time series. Irregular means that the time between consecutive time points can vary. Given two timestamps t_1_ and t_2_, Bahadori and Liu define a Gaussian kernel function

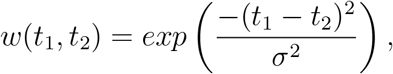

where σ represents the effective kernel width. Based on this kernel function, the operator ⊙ defined below generalizes the inner product for two ‘irregular’ time series — *x*, sampled at times *t_x_*(1),*t_x_*(2),…, *t_x_*(*N_x_*), and *y*, sampled at times *t_y_*(1), *t_y_*(2),…, *t_y_*(*N_y_*) — as

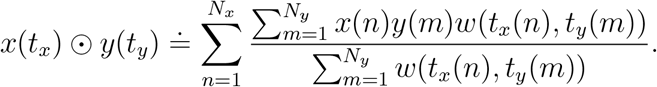

*N_x_* and *N_y_* can differ, which SINGE exploits for its dropout handling and subsampling (Section 3.2).

We now have *P* irregular time series *x*_1_, *x*_2_, …, *x_P_* obtained from a linear and stationary VAR process. Each series *x_i_* of length *N_i_* is sampled at irregularly-spaced timestamps *t_i_* such that *t_i_*(*n* +1) ≥ *t_i_*(*n*) for *n* = 1, 2,…, *N_i_* — 1. As with the Lasso Granger Causality test, the objective of the GLG test is to obtain the sparse coefficient matrix a_*i*_, which represents the underlying VAR model for the target series x¿. To overcome the irregularity of the time series, we follow Bahadori and Liu by visualizing each vector a_*i,j*_ of the coefficient matrix as a time series 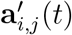 with respect to a given timestamp *t*. This is accomplished by assigning a new timestamp *t_a_*(*l*) = *t — l*Δ(*t*) to each lagged coefficient *a_i,j_*(*l*), where Δ*t* represents the time-lag between successive lagged coefficients in *a_i,j_*. Thus, this time series can be viewed as the sequence of (lagged time, coefficient value) pairs.

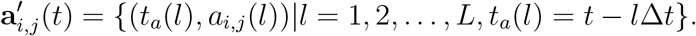

Note that for *t*_1_ = *t*_2_, the two time series 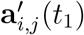 and 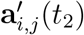 would have the same coefficient values *a_i,j_* (*l*) but different lagged timestamps. For example, if we select hyperparameter values Δ(*t*) = 5, and *L* = 3, then, for *t*_1_ = 50 and *t*_2_ = 75, we have

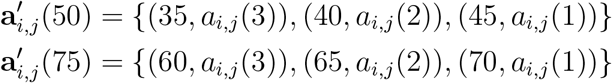

respectively.

Next, for a given timestamp *t_i_*(*n*) corresponding to a sample in *x_i_*, we generalize the inner product in Equation 2 by using

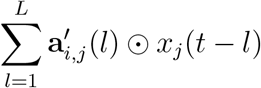

defined on 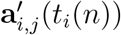 and *x_j_* (*t_j_*) using their respective timestamps to calculate the kernel weights. Substituting this generalized inner product in Equation 2, we obtain the optimization problem for GLG, given by

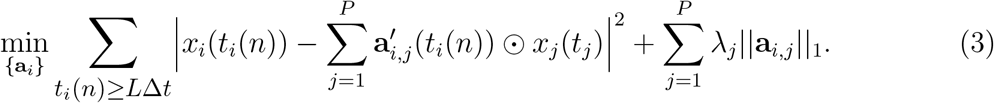

The first term represents the mean-squared error between the sample values *x_i_*(*t_i_*(*n*)) of the *i*-th series at each timestamp *t_i_*(*n*) ≥ *L*Δ*t* and its corresponding prediction from the generalized inner product 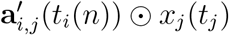, which uses the kernel defined above to ‘smooth over’ the mismatched irregular timestamps. The second term is a sparsity constraint on the coefficient matrix a_*i*_. The minimizer a_*i*_ of the objective function in Equation 3 provides the coefficient matrix that represents the VAR model of the target series *x_i_* from all available source time series *x_j_*, with λ determining the sparsity of the coefficient matrix. If the time series represent irregularly-spaced gene expression data, a_*i*_ can be interpreted as an estimate of the regulatory effect of other genes on the *i*-th gene. The presence of edges in the regulatory network for the *i*-th gene is indicated by significant non-zero values in the matrix a_*i*_. The ‘edge weight’ of *j → i* can be quantified by ||**a**_*i,j*_||_2_, ||**a**_*i,j*_||_∞_, or |∑_*l*_a_*i,j*_(*l*)|, the latter aiming to capture the net impact of gene *j* on gene *i*. In the default GLG setup, the individual weights in the *l*_1_-constraint of the above equation are assigned the same value, with λ_*j*_ = λ. However, because we are not interested in the auto-regulation of *x_i_*, we remove the sparsity constraint on the autoregressive edge (λ_*i*_ = 0) in order to reduce the number of false positives in the cross-regulatory relationships, where sparsity is typically enforced with a positive λ_*j≠i*_ = λ.

The optimization problem in Equation 3 can be solved P separate times to infer the regulators of all P genes in the network. The GLG-identified regulators are obtained as the smallest group of genes whose past expression values are most predictive of gene i’s time series expression values. Because the core algorithm of the GLG test is implemented using the *glmnet* package [99], it supports count-based expression data (e.g. from unique molecular identifiers) by assuming a Poisson distribution for the expression levels.

### 3.2. Single-cell Inference of Networks using Granger Ensembles

In this section, we describe how the SINGE algorithm, which has the GLG test at its core, infers GRNs from single-cell expression data. The SINGE algorithm takes ordered single-cell RNA-seq data as input, with an optional zero-handling pre-processing step to mitigate the effect of dropouts. The data are analyzed using multiple GLG instances with different hyperparameters, each inferring possibly differently ranked regulator-gene interactions. These ranked inferences are aggregated using a modified Borda count, with an optional subsampling stage increasing the effective ensemble size.

#### 3.2.1. SINGE Input

*Ordered Single-Cell Gene Expression Data.* The input to SINGE is ordered single-cell gene expression data, with a pseudotime assigned to each cell that represents its position along the biological process. Given ordered single-cell data, the pseudotimes are first normalized to a scale of 0-100. Thus, the first cell represents 0% progress, and the last cell represents 100% progress through the biological process. The distribution of cells’ pseudotimes is not uniform. As a result, each gene’s expression data is an irregularly-spaced time series in the pseudotemporal reference. We represent each gene’s expression trend along the pseudotemporal reference as an augmented series with both the pseudotimes and the gene expression values. That is, for the i-th gene, we create the series (*t_i_, x_i_*), where *x_i_* is the time-series representing the gene’s expression and t¿ represents the pseudotime of the corresponding cell. As of version 0.5.0, SINGE is applicable to all acyclic trajectories.

*Ordering Cells via Trajectory Inference.* If the single-cell dataset is not already ordered, any cell ordering method that assigns continuous pseudotimes (Section 1) can be used to annotate the cells before running SINGE. For the retinoic acid-driven differentiation dataset, we apply Monocle 2 [25], which uses reverse graph embedding to identify branching processes. For other trajectory inference methods, we use the `dynverse` package [18], which provides a streamlined approach to benchmark and use trajectory inference algorithms and can shortlist algorithms based on trajectory type, dataset size, and other criteria.

*Regulator Indices (Optional).* By default, SINGE assumes that all genes are potential regulators. It can optionally limit the possible regulators using regulator indices regix ⊆ {1, 2,…, *P*} provided by the user. SINGE will infer a GRN with potential regulator genes limited to only those corresponding to the regix indices. Using regulator indices can substantially improve SINGE’s runtime.

*Branching Information (Optional).* By default, SINGE assumes that the ordered singlecell data correspond to a linear biological process. However, if the inferred trajectory has a branching topology, this information can be passed to SINGE by creating an optional binary matrix called branches with *N_ceus_* rows and *N_branches_* columns. If *branches*[*i, b*] = 1, it represents the membership of the *i*-th cell in the b-th branch of the trajectory. This allows SINGE to handle any type of acyclic trajectory.

#### 3.2.2. Zero (Dropout) Handling

One of the most prominent technical artifacts in single-cell RNA-seq is dropout. This is manifested as a large number of zero readings due to inefficiencies in mRNA capture in the measurement process. Dropout causes the measured expression data to contain a higher number of zeros than the true biological zeros [100]. There have been efforts to overcome this problem by imputing the missing values [100, 101]. However, inappropriate imputation can negatively impact differential expression testing [102] and can have a positive, neutral, or negative effect on Monocle’s pseudotimes depending on the choice of algorithm [103].

If we remove the zero-valued measurements altogether from the dataset, GLG effectively imputes the missing values without an external imputation algorithm by virtue of its kernel-based approach for analyzing irregular time series. Thus, depending on the severity of the dropout, SINGE contains an optional step of removing some of the zeros and the corresponding pseudotime values. This can be achieved through an additional hyperparameter *prob-zero-removal*. For each gene, each zero-valued sample and its corresponding pseudotime are removed with probability *prob-zero-removal*.

#### 3.2.3. Hyperparameter Diversity

The primary hyperparameters in the GLG tests include the sparsity constraint λ, the time resolution Δt between the elements of the vector **a**_*i,j*_, the length L of the vector a_*i,j*_ (which determines the extent of the lagged time series for the GLG analysis), and the kernel width σ. The zero-handling stage introduces another optional hyperparameter *prob-zero- removal*.

If the process being studied is a stationary process containing simplistic regulatory networks, the above hyperparameters could potentially be tuned to optimize cross-validation performance. However, transcriptional regulation is non-linear and non-stationary in nature. A single GLG test, however optimal its settings, can produce false positives due to the assumption of linear and stationary causal relationships. In addition, there may not be a single set of hyperparameters that are optimal for all regulatory interactions. To overcome this, we analyze the data using multiple GLG tests with diverse hyperparameters and aggregate the rankings obtained from the individual GLG tests (Section 3.2.5). Our assumption is that the top-ranked regulatory edges that consistently appear for many hyperparameter combinations are enriched for true positive interactions.

#### 3.2.4. Subsampling Stage

SINGE includes an optional stage that increases the effective ensemble size by subsampling versions of the original single-cell data. The subsampling can make the inferred GRN more robust to outliers in the gene expression data. Specifically, for each hyperparameter combination, we generate *N_subsample_* (default 10) data replicates. Because GLG can handle irregular time series, we have the option to use two different strategies for subsampling. The simplest strategy would be to randomly remove a small subset of cells from the dataset. This ensures that all genes’ pseudotime series have the same pseudo-timestamps. However, removing entire cells could ignore important cells in rare transient states.

We instead use an alternate strategy that randomly removes samples independently from each gene’s pseudotime series. Using this strategy, the probability of removing an entire pseudo-timestamp (cell) is greatly diminished. However, no two genes have the same series of pseudo-timestamps, each has a unique irregular time series with high probability. In our experiments, we independently remove samples for each gene using a probability of sample removal of 0.2, the SINGE default. The SINGE subsampling is similar to bagging [104] except that the sampling is without replacement and it uses a different aggregation approach.

#### 3.2.5. GLG Runs and Modified Borda Aggregation

After enumerating all hyperparameter combinations and subsampled replicates, SINGE runs GLG on each subsampled replicate using the different hyperparameter combinations. At the end of each GLG test, we obtain an adjacency matrix **A** using

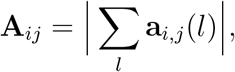

where **a** is the *P × P × L* coefficient matrix output from the GLG test. The matrix **A** represents one candidate GRN, with the magnitude of each element representing the edge weight assigned to the corresponding regulator-gene interaction. These edge weights are used to rank the possible regulator-gene interactions. A rank is assigned to only those interactions that correspond to a nonzero element of **A**.

Once the rankings from the GLG tests on all hyperparameter combinations and subsampled replicates are obtained, we aggregate them using a modification of the Borda count [105]. The Borda count aggregates ranked lists by defining a scoring rule that assigns weights to the items in each ranked list and summing the weights to obtain a final consensus ranking. The goal is to favor items, in our case regulator-gene interactions, that are consistently ranked high over those that are ranked high only occasionally or not at all. The traditional Borda count scoring rule assigns a weight of *N* for the first ranked item in a list, *N* — 1 to the second, and so on [105]. Alternative scoring rules, such as the Dowdall rule [106], assign weights that decay more quickly, placing more relative importance on the top-ranked items.

We use a scoring rule that assigns weights of 1/*i*^2^ for the *i*-th ranked interaction within each individual ranked list from a single GLG test. The weight is zero for an unranked regulator-gene interaction. This scoring rule was selected based on empirical tests with the ESC to endoderm differentiation dataset. The final SINGE score of each interaction is obtained by summing the weights assigned to that interaction across all ranked lists. This score is subsequently used for the final GRN edge ranking. We also obtain the top regulators of the biological process by summing the SINGE scores of all outgoing edges for each regulator and sorting the regulators in order of decreasing magnitude. In the case of branching processes, SINGE’s default behavior is to perform the modified Borda aggregation on all branches together to obtain one output for the overall branching process. Alternatively, a user can obtain branch-specific SINGE outputs by storing the individual GLG test results in separate branch-specific directory and performing the modified Borda aggregation on each set of results separately.

Similar ensembling and aggregation strategies are widely used in GRN inference in order to improve the robustness of the predicted networks, reduce sensitivity to noise, and avoid false positives. SINGE’s modified Borda count aggregation is one specific strategy among many related ideas. It emphasizes the interaction ranking instead of the magnitude of the **A**_*ij*_ coefficients, which are difficult to compare directly when combining results from GLG runs that use different degrees of regularization λ. SINGE’s aggregation is closely related to the stability selection [107] in TIGRESS [108] except SINGE aggregates predictions over many hyperparameter combinations and its randomization comes from the randomly removed observations during the subsampling stage instead of randomly rescaling TF expression. Unlike other unsupervised aggregation approaches [109], SINGE’s modified Borda counts do not assume that the ranked interaction lists are conditionally independent. Furthermore, SINGE’s aggregation does not require generating a null distribution of **A**_*ij*_ coefficients from permuted data [46], which is computationally more expensive but has the benefit of providing interaction false discovery rates.

#### 3.2.6. Case Study Hyperparameters

Table 3 lists the hyperparameter values used to generate the GLG ensembles for all case studies. The subsampling stage creates 10 replicates for each hyperparameter setting by removing samples from individual gene expression values with probability of sample removal 0.2. Thus, not only is each time series irregular, but it has partially different time references compared to the other time series in the data set. For most of the main case studies, we use the default mode with *prob-zero-removal* = 0, only changing it when we analyze its effect on GRN performance (Figure 9) or to reduce runtime on the bone marrow dataset (Section 3.3.3). The total number of GLG tests, accounting for hyperparameter diversity and subsampling, is

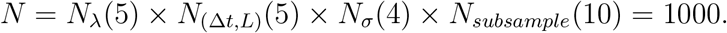

**Table 3:** Hyperparameter combinations considered. λ is the sparsity parameter, Δ*t* determines the time resolution, *L* represents the number of time lags under consideration, and *σ* represents the kernel width used for GLG. Only specific pairs of Δ*t* and *L* are considered instead of all possible combinations.

| Hyperparameter(s) | Values |
| --- | --- |
| $\lambda$ | 0, 0.01, 0.02, 0.05, 0.1 |
| $(\Delta t, L)$ | (3, 5); (5, 9); (9, 5); (5, 15); (15, 5) |
| $\sigma$ | 0.5, 1, 2, 4 |

The subsampling approach and the consensus-rewarding nature of the modified Borda aggregation stage reduces the need to optimize the hyperparameter combinations for each dataset. The outputs from GLG tests on each subsampled replicate will better agree with each other for hyperparameters that are suitable for the dataset.

### 3.3. Datasets

#### 3.3.1. ESC to Endoderm Differentiation

The first dataset is from Hayashi et al. [51], who collected single-cell RNA-seq data from 456 cells at five time points over a 72 hour duration in which primitive endoderm cells were differentiated from mouse embryonic stem cells. Matsumoto et al. [15] used Monocle [25] to order these cells along the differentiating process, assigning a pseudotime to each cell. We use their Monocle results in our analyses. The expression dataset is limited to 100 TFs exhibiting the highest variance in expression value and 356 cells.

#### 3.3.2. Retinoic Acid-driven Differentiation

The second dataset was obtained from Semrau et al. [54], where SCRB-seq data was collected at nine times during a 96 hour period from mouse embryonic stem cells differentiating into neuroectoderm and extraembryonic endoderm-like cells. We order the cells using Monocle 2 [25], with the ordering genes chosen by Monocle 2 in an unsupervised manner by identifying genes that are differentially expressed in response to the introduction of the growth medium. Although Matsumoto et al. [15] applied the original Monocle to the first dataset and we retain their pseudotimes, we prefer Monocle 2 for this case study, the most recent version available at the time of the analysis. After ordering, we limit the scope of the analysis to the 1886 cells along the longest trajectory of the differentiation process (Figure S1) exhibiting non-trivial expression levels. Once Monocle 2 orders the cells along a pseudotemporal reference, it can find genes that change in expression as cells progress along pseudotime. We retain the top 626 differentially expressed genes (*q* < 10^-5^) along the pseudotime ranked by Monocle 2 for testing the GRN algorithms.

To obtain another version of the retinoic acid-driven differentiation dataset, we use `dynverse` to select an appropriate alternative trajectory inference method. Based on the expected branching topology of the trajectory, we use the graphical user interface `dynguidelines` to narrow our search to the top four recommended methods: Slingshot [110], SCORPIUS [111], PAGA, and PAGA Tree [95]. Running trajectory inference on the entire dataset does not yield any meaningful branching trajectories for any these methods. When we instead limit the genes to the same 626 differentially expressed genes from our Monocle 2 analysis, we obtain a reasonable branching trajectory with PAGA Tree (Figure S11). The `dynverse` wrapper for the PAGA Tree algorithm is version 0.9.9.06. We choose the longest branch (backbone) of this inferred trajectory as an alternate input for SINGE network inference.

#### 3.3.3. Mouse Bone Marrow Mesenchyme to Erythrocyte Differentiation

Starting with the dataset available from the Mouse Cell Atlas [87] at https://figshare.com/s/865e694ad06d5857db4b, we use the `dynverse` utility to select an appropriate trajectory inference method. Based on the expected topology of the trajectory, we use the graphical user interface `dynguidelines` to prioritize four appropriate trajectory inference methods: Slingshot [110], Embeddr [88], SCORPIUS [111], and PAGA [95]. Of these, the Embeddr inferred trajectory was most consistent with the reference trajectory provided in the `dynverse` database. The `dynverse` wrapper version used for Embeddr is 0.9.9.01, and the resulting trajectory is shown in Figure S12. The ordered single-cell dataset has 3025 genes and 3105 cells in a linear trajectory. We normalize the count-based expression data using a log(x+1) transformation [112].^1^ Finally, because the dataset is large but sparse, we use *prob-zero- removal* = 0.75, which makes the regression problem smaller by dropping many zero-valued samples and speeds up SINGE.

#### 3.3.4. dyngen *Synthetic Data*

We generate two simulated datasets using the dyngen package [93] available at https://github.com/dynverse/dyngen (git commit 73192c82f39e1b4318aea56d4c87ed02c1bf145e). Both datasets come from the same GRN that has 140 genes — 25 TFs, 15 housekeeping genes, and 100 target genes — and 164 edges, most of them originating from TFs. The first dataset contains 1000 cells, and the second contains 20000 cells. The script used for generating the datasets, the datasets in SINGE input format, and the gold standard GRN are provided at https://github.com/gitter-lab/SINGE-supplemental and archived on Zenodo (https://doi.org/10.5281/zenodo.3627325). The 20000 cell version was used solely to evaluate the computational runtime for larger datasets.

### 3.4. Evaluation

To evaluate the GRNs from the ESC to endoderm differentiation and retinoic acid-driven differentiation datasets, we use the ESCAPE database [52] as a gold standard, namely the cataloged ChIP-chip, ChIP-seq, loss-of-function (*lof*), and gain-of-function (*gof*) experiments. Each GRN method ranks the possible edges in the network in order of confidence. We plot the respective precision-recall curves and compute the average precision (**A**) and average early precision (**E**) for comparison.

We evaluate the GRNs from the simulated dyngen dataset using the dyngen model as a gold standard. This simulated GRN only contains direct edges from regulators to targets. We also evaluate the predicted networks with respect to extended gold standards that include indirect interactions, which considers transitive interactions. That is, if gene interaction *A → B* and *B → C* exist, whether direct or indirect, then *A → C* also exists in the modified gold standard. Starting with the original direct `dyngen` GRN with 164 interactions, we iteratively apply this transitive property to obtain gold standards with different levels of indirect gene interactions. These indirect networks contain 352 edges after the first iteration, 744 edges after the second iteration, and 921 edges after the third iteration. Further iterations do not expand the gold standard GRN.

#### 3.4.1. ESCAPE Database

The ESCAPE database [52] is a repository cataloging numerous experiments conducted on human and mouse ESCs. We use the ChIP-chip, ChIP-seq, *lof*, and *gof* experiments as a gold standard to evaluate the inferred GRNs. Despite ESCAPE being one of the most comprehensive repositories of such experimental results, it does not have reference data for all predicted regulators. Therefore, we evaluate the inferred networks using the sub-matrix for which the gold standard is available.

To generate the gold standard, we combine all gene interactions in the ChIP-chip/ChIP-seq and *lof/gof* databases related to the genes from the single-cell data being analyzed. Gene interactions not documented in the ESCAPE databases are assumed to not exist. However, this approach can lead to a high number of false zeros in the gold standard if a particular regulator was not studied genome-wide. For example, whereas ESCAPE documents thousands of ChIP-chip/ChIP-seq interactions for most TFs, two of the TFs report less than 200 interactions. To avoid false zeros in the gold standard, we generate our gold standard using only regulators with at least 1000 target genes in the ChIP-chip/ChIP-seq data and 500 target genes in the *lof/gof* data.

#### 3.4.2. Average Early Precision

Because a majority of SINGE’s hyperparameter sets predict a sparse regulatory network, it is better suited to ranking the top GRN interactions instead of ranking all of them. Average precision, which considers the entire precision-recall curve, may not be the ideal performance metric for evaluating such methods. In addition, the top-ranked regulator-gene interactions are the most relevant for prioritizing experimental studies. Therefore, we also consider the average early precision, which evaluates the inferred network by calculating the average precision up to a partial recall threshold. We use a partial recall threshold of 0.1. That is, average early precision evaluates the ranking performance of GRN inference methods up to the point where they identify 10% of known gene interactions according to the gold standard. For consistency across the evaluations, we generally assess whether there is a 10% improvement in average precision or average early precision when comparing two GRNs. If the performance of one GRN is not at least 10% better than the other, we consider them to be approximately similar.

#### 3.4.3. Existing GRN Methods

In addition to SINGE, we use GENIE3, SCODE, SINCERITIES, and Jump3 to infer GRNs. Originally intended for use with bulk transcriptomics, the GENIE3 algorithm [12] can also be applied to single-cell data [113] and acts as a reference method that does not use pseudotemporal ordering information. We installed GENIE3 version 1.6.0 from BioConductor https://bioconductor.org/packages/release/bioc/html/GENIE3.html and used its default operating mode. We downloaded SCODE from the GitHub repository https://github.com/hmatsu1226/SCODE (git commit 28acad67893c0fba7eeee670c339809d 45ae6377) and used the same settings as in Matsumoto et al. [15] for the ESC to endoderm differentiation dataset with D = 4 degrees of freedom in the expression dynamics. We used *D* = 20 for the retinoic acid-driven differentiation dataset to account for the much larger network of 626 genes. We obtained the SINCERITIES toolbox dated 16 December 2016 from http://www.cabsel.ethz.ch/tools/sincerities.html and used the default settings. An equivalent version of the Jump3 code we used can be obtained from https://github.com/vahuynh/Jump3 (git commit 03a7e86d82f2383c56fd11c658dfce574fbf1a1a). In contrast to the other methods, Jump3 uses only ordering information. We used *noiseVar.obsnoise* = 0.1, but all other settings were the defaults. Because Jump3 did not terminate in a reasonable amount of time on the full retinoic acid-driven differentiation dataset, we reduced the dataset by arbitrarily dropping cells with probability 0.5. Despite this reduction in the data size, the Jump3 algorithm did not converge for two target genes, Tdh and Vdac1. As a result, we rank the corresponding edges at the bottom of the ranked list, which could affect the quality of the Jump3 results for the retinoic acid-driven differentiation dataset.

#### 3.4.4. KinderMiner and Gene Ontology Enrichment

We performed KinderMiner (v1.5.4) [75] analysis on the SINGE top 20 regulators to search for known associations of these genes with the three keyphrases ‘embryonic stem cells,’ ‘neural development,’ and ‘endoderm development’ in a local collection of 26877474 PubMed abstracts downloaded from NCBI in December 2018. We report the statistically significant associations (p < 10^-4^) in Table 1 using the labels ‘ESC,’ ‘NeurDev,’ and ‘EndoDev,’ respectively. The significance threshold corresponds to a family-wise error rate of *FWER* < 6 × 10^-3^, accounting for a family size of 60 gene-keyphrase pairs. In Supplementary File 4, we provide the raw KinderMiner results obtained using the search setting **anySpeciesSEP**. This corresponds to a species agnostic search in which words of keyphrase can be anywhere in the PubMed abstract.

We also performed functional profiling of the ordered 626-gene list from SINGE using the g:GOSt tool in g:Profiler [55] version r1760_e93_eg40. We consider only Gene Ontology [114] biological process terms and specify ‘mus musculus’ as the organism. The candidate regulator list from SINGE is ordered, so we use the ‘ordered query’ option, which allows g:Profiler to perform incremental enrichment analysis over the gene list. The significance threshold used was Fisher’s one-tailed test, the default test for g:GOSt, with multiple testing correction using the default g:SCS method. Supplementary File 3 provides the complete output of the g:GOSt test. The significance test considers the entire ranked regulator list, but we highlight only the top 20 regulators in Table 1. In addition, we derived the loss-of-function phenotypes in Table 1 from the Mouse Genome Databases Mammalian Phenotype Ontology Annotations [115].

#### 3.4.5. SINGE Software Availability and Versions

A MATLAB implementation of SINGE is available at https://github.com/gitter-lab/ SINGE under the MIT license and archived on Zenodo (https://doi.org/10.5281/zenodo.2549817). The GitHub repository contains the datasets and default hyperparameter settings used in the manuscript, scripts to generate custom hyperparameter files, and compiled code so that SINGE can be run without a MATLAB license. We also include a Docker image (https://hub.docker.com/r/agitter/singe). Supplemental scripts and analysis are available at https://github.com/gitter-lab/SINGE-supplemental and archived on Zenodo (https://doi.org/10.5281/zenodo.3627325).

We used SINGE version 0.1.0 for nearly all analyses, except Figure 11 and Table 2, which also include results from newer versions of SINGE as indicated. In addition, the analyses in Sections 2.2.4 and 2.2.5 were performed using only version 0.3.0, which included code optimizations for improving stability and compute time for large datasets such as those encountered in the aforementioned sections. See https://github.com/gitter-lab/SINGE for the latest release notes and usage recommendations.

## 4. Discussion

SINGE is a GRN reconstruction algorithm that adapts Granger Causality to detect dependencies in single-cell gene expression data annotated with pseudotimes. Although it was designed for single-cell data, the kernel-based smoothing could also be valuable for bulk time series gene expression data when the time points are irregularly spaced or individual expression samples are noisy. SINGE has the potential to prioritize regulators for future DNA-binding or functional studies. For example, in the retinoic acid-driven differentiation study, many of the top-ranked SINGE regulators (Table 1) are enriched for relevant differentiation process and regulatory annotations but have not yet been characterized in the ESCAPE database.

When assessed in the retinoic acid-driven differentiation case study, in which none of the GRN methods’ settings were tuned to optimize performance on this dataset, SINGE has better precision-recall performance than four existing methods. However, we caution that single metrics like average precision can be misleading. Closer inspection reveals that SINGE’s better-than-random precision-recall performance in Figure 5 is driven by its ability to identify important regulators and assign them a higher rank, as seen in Figure 7. In contrast, Jump3’s better-than-random precision-recall performance in Figure 5 is driven by its better-than-random performance for many individual regulators, as seen in Figure 6. Because the precision-recall curve for the entire GRN can mask near-random performance for many individual regulators, we recommend regulator-specific visualizations (Figures 6 and 7) to provide more context.

We designed SINGE for a high-throughput computing environment, ensembling many GLG tests under different hyperparameters and using data subsampling to improve robustness and performance. This approach makes SINGE more resilient to dropout in the single-cell gene expression data and less sensitive to the hyperparameter ranges tested. Ensembling strategies have proven effective in a variety of GRN inference settings, such as DREAM challenges [8]. Our use of modified Borda aggregation for ensembling emphasizes the top-ranked, most-confident predictions.

### 4.1. Caveats and Limitations

Granger Causality has a precise statistical meaning built upon the idea that causes must come before their effects [40]. It is useful in practice but does not necessarily satisfy more general definitions of causality [40]. The main assumption we make by using GLG is that the expression data are obtained from a linear and stationary VAR model. However, complex biological systems have dynamic, non-linear gene interactions and are expected to generate non-stationary expression trends. Violating the assumptions of linearity and stationarity can have a significant impact on the performance of individual GLG tests. Furthermore, Granger Causality tests result in false positives in scenarios with hidden variables [116]. However, these discrepancies between theory and practice are commonly accepted in biological applications of Granger Causality [43, 117]. In addition, SINGE’s Borda aggregation helps to push the most robust edges in the network to the top of the final ranked list.

Some of the Granger Causality-related drawbacks potentially could be addressed by integrating SINGE with complementary data types. The relationship between TF concentration and transcriptional activity represents only one type of transcriptional dynamics, neglecting epigenomic modifications, TF post-translational modifications, TF localization, and transcriptional co-factors [118]. GRN inference can be more accurate when using ChIP-chip, ChIP-seq, protein-protein interactions, regulator *lof/gof* experiments, or DNA binding motifs as prior knowledge on the network structure [119, 120] (reviewed in Chasman et al. [9]). Other single-cell GRN inference algorithms have incorporated priors. SOMatic [121] and Symphony [122] use single-cell ATAC-seq and scdiff [123] integrates TF-gene interactions. To model prior information in SINGE, we could assign different penalty factors λ_*j*_ for the *j*-th regulator of target gene i based on the prior probability of the edge *p_ij_*. An alternative would be to use SINGE output in conjunction with the supplementary sources of information and aggregate all the information after-the-fact [124, 125]. However, the current version of SINGE intentionally uses only gene expression data because integrative approaches can benefit from understanding the best ways to infer GRNs from expression data alone. This also makes SINGE widely applicable in conditions and species where suitable priors are not available.

Another assumption of SINGE, SINCERITIES, and SCODE is that the pseudotime values assigned to individual cells have a high fidelity. However, in the retinoic acid-driven differentiation dataset, SINGE performs better when the Monocle 2 pseudotime values are disregarded and only the cell ordering is used (Figure 10). On the other hand, using PAGA Tree pseudotimes improves the precision-recall performance when compared to the Monocle 2 pseudotimes. GRN performance in SINGE and related methods depends on the quality of the pseudotime values. Assigning uninformative pseudotime values to ordered cells can be more detrimental to the network inference performance than simply using the order without pseudotimes.

We propose that pseudotimes’ impact on GRN accuracy could be used to evaluate pseudotime inference algorithms, complementing other benchmarking metrics for pseudotimes [18]. Integrating GRN benchmarking [126] and `dynverse`’s pseudotime benchmarking [18] would enable us to systematically evaluate which types of pseudotimes best support network inference and empirically assess the types of GRN motifs that cannot be unambiguously recovered from single-cell expression data [127]. In addition, Qiu et al. [33] proposed that RNA velocity [128] may help overcome limitations of pseudotime for GRN reconstruction.

### 4.2. Benchmarking and Evaluation

Inferring gene regulatory networks only from single-cell gene expression data is a difficult task. One evaluation of network inference algorithms on simulated single-cell datasets reported that their performances were only slightly better than a random edge ordering [129]. New single-cell gene expression simulators designed specifically for simulating GRNs [126, 130] can help inform which GRN inference methods are best for different types of biological trajectories. Both of these studies evaluated SINGE performance with their GRN simulators but with different ensembling strategies. BEELINE [126] optimized SINGE’s hyperparameters and constructed much smaller ensembles than we do in this study. SERGIO’s ensembling [130] was much closer to ours except for differences in the treatment of the *λ* hyperparameter. In the SERGIO evaluation, SINGE was the best GRN method when inferring networks from simulated datasets with added technical noise.

An important aspect when evaluating network inference on experimental data is the relevance of the gold standard. In the SCODE evaluation [15], the gold standard was TF binding interactions estimated from DNaseI footprints and sequence motifs. However, it was merged across all human and mouse cell types instead of only those relevant to the mouse ESC to endoderm differentiation process. In the Chen and Mar benchmarking of stem cell datasets [129], the gold standard consisted of all interactions from the STRING database [131]. These included interaction types that are not directly informative about transcriptional regulation and were not limited to the specific cell types of interest. For our evaluation, we limit the gold standard to data from mouse embryonic stem cells obtained from ChIP-chip/ChIP-seq and *lof/gof* studies cataloged by ESCAPE [52], which are more relevant to the biological processes we study and more directly indicative of transcriptional relationships. BEELINE showed that the choice of a cell type-specific versus non-specific gold standard has a substantial impact on GRN inference performance metrics [126].

There remain open questions regarding appropriate evaluation methodologies. For example, we combine ChIP-chip/ChIP-seq and *lof/gof* information, but the precision-recall performance of the GRN methods is quite different when examining ChIP-chip/ChIP-seq or *lof/gof* data alone (Figure S3). These two types of data are known to have low overlap [132]. Our evaluation suggests the SINGE’s search for lagged gene expression dependencies may detect more indirect regulatory relationships than direct TF binding, which is consistent with our `dyngen` evaluation. We observe SINGE’s precision-recall performance improves relative to the the baseline precision when adding indirect gene interactions to the `dyngen` gold standard.

Although the GRN methods we evaluate have better than random average precision when assessing the entire network, they are only marginally better than random when ranking outgoing edges from individual regulators. For the regulator-specific average early precision, each GRN method is better than random for only some regulators. Precision-recall is preferable to the receiver operating characteristic for evaluating biological network inference due to the sparsity of the gold standard [133], but the average precision for the entire network may overestimate the utility of GRN inference methods for studying individual regulators (Figures 5 and 6).

### 4.3. Future Work and Extensions

The current version of SINGE can handle all biological processes with acyclic trajectories. However, SINGE provides just one regulatory network for the entire trajectory. In the future, we could add functionality to identify branch-specific networks and interactions. An alternative approach would be to adapt SINGE to treat each branch as a task in a multitask GRN inference problem [134]. In addition, the kernel could be modified so that certain pseudotime intervals can be considered more informative, for example, the interval around a major bifurcation point.

Because each pseudotime estimation method orders the cells based on different assumptions and algorithms, no two methods will result in the same cell ordering. Some trajectories are better suited for a particular biological process, but we cannot objectively verify the correctness of the trajectory and cell ordering. To mitigate the effect of the pseudotime estimation method on network inference, we could integrate SINGE with the `dynverse` framework and expand the ensemble to include GLG test results from multiple pseudotime estimation methods.

SINGE uses a common *prob-zero-removal* for all genes, but the algorithm can be modified to incorporate gene-specific zero removal probabilities. Future work may involve a more sophisticated zero-handling approach, which would remove only those zeros that are inconsistent with other non-zero measurements from similar cells. Methods like SCONE [94] can provide additional information for removing zeros more selectively than the current approach.

Other elements of the GLG regression framework can be adapted as well. Nguyen and Braun [135] place a monotonicity constraint on the coefficients **a**_*i,j*_ such that the more recent coefficients have higher magnitude than the more distant ones. Similarly, we could adapt the kernel to give higher weight to more recent samples in the pseudotime than more distant ones. Another possible direction involves exploration of the kernel-based generalizations to the Group Lasso [136, 137]. This would enable SINGE to regularize all coefficients from a regulator as a group instead of treating different lagged coefficients as separate variables. In general, the kernel-based approach at the core of SINGE provides great flexibility to adapt our GRN reconstruction algorithm to emphasize different aspects of dynamic biological processes.

## Acknowledgements

We thank Christina Kendziorski for feedback on the SINGE algorithm, Alireza Fotuhi Siahpirani for assistance with the ESCAPE database, Rafael Feliciano for discussion of the GRN evaluation, Scott Swanson for testing the SINGE software, Robrecht Cannoodt and Wouter Saelens for help with the `dyngen` simulation, Eric Bolden for support compiling SINGE for macOS, and members of the Gitter, Stewart, and Kendziorski research groups for their helpful comments.

## Supplementary Files

- Supplementary Information containing Figures S1 to S12
- Supplementary File 1. Predicted GRNs on the ESC to endoderm differentiation dataset (Figure 3)
- Supplementary File 2. Predicted GRNs on the retinoic acid-driven differentiation dataset (Figures 5 and 11)
- Supplementary File 3. List of genes ordered according to SINGE influence for the retinoic acid-driven differentiation dataset with corresponding g:Profiler enrichment results
- Supplementary File 4. KinderMiner associations for the top 20 regulators according to SINGE influence for the retinoic acid-driven differentiation dataset
- Supplementary File 5. Predicted GRN from SINGE on the mouse bone marrow dataset, including the top 10000 SINGE ranked edges and list of regulators ordered according to their SINGE influence.

## Supplementary Information

**Figure S1:**
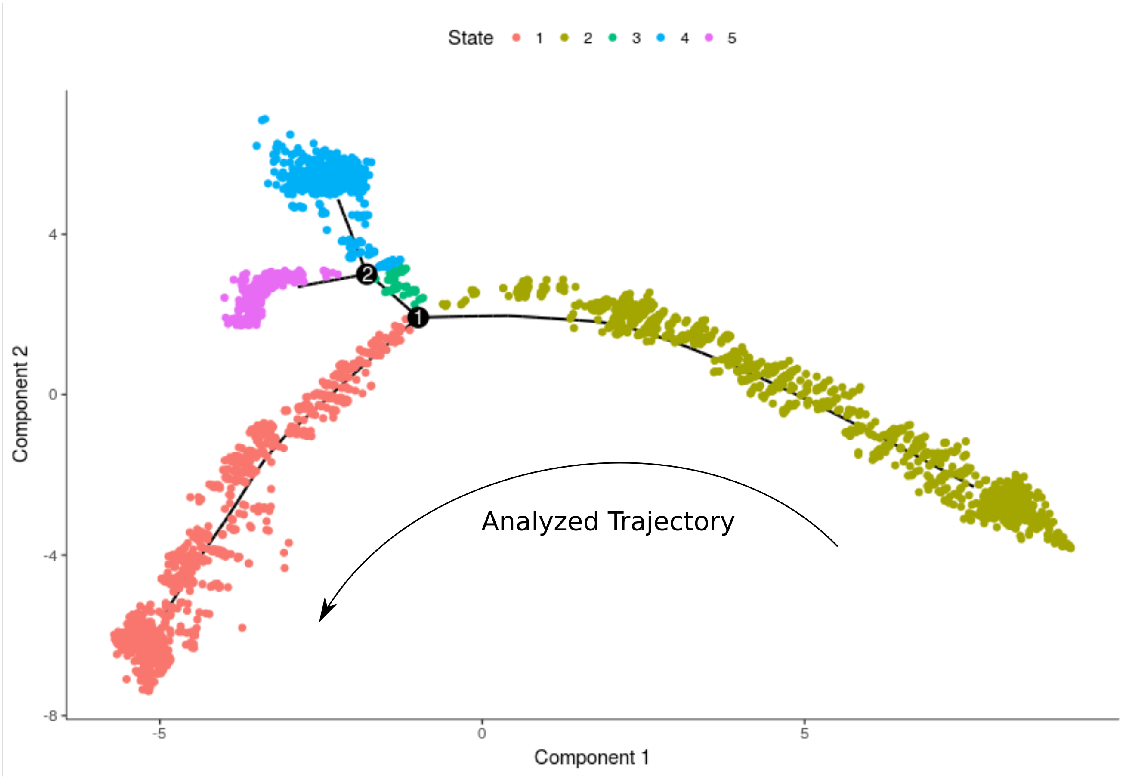
Monocle 2 trajectory of the retinoic acid-driven differentiation process. We analyzed the trajectory constituting states 2 (early part comprising mostly data collected at 0h) and 1 (later part including cells collected at 96h).

**Figure S2:**
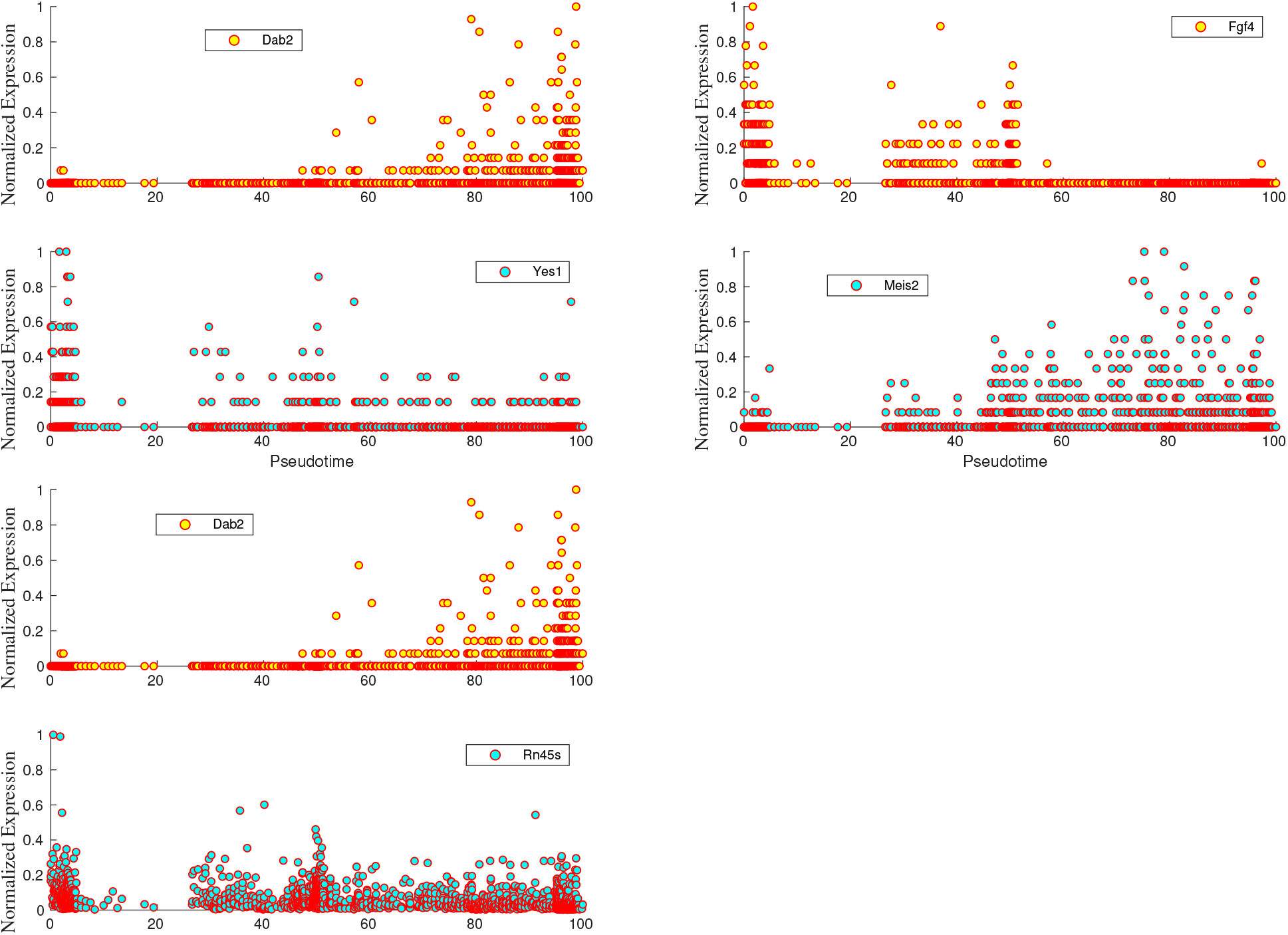
Expression trends of genes from three of the top-ranked SINGE interactions (Dab2→Yes1, Fgf4→Meis2, and Dab2→Rn45s) in the retinoic acid-driven differentiation dataset. From the expression trends, we see a negative correlation between Dab2 and Yes1, as well as between Fgf4 and Meis2. The relationship between Dab2 and Rn45s is not obvious from the expression trends alone and is likely a false positive.

**Figure S3:**
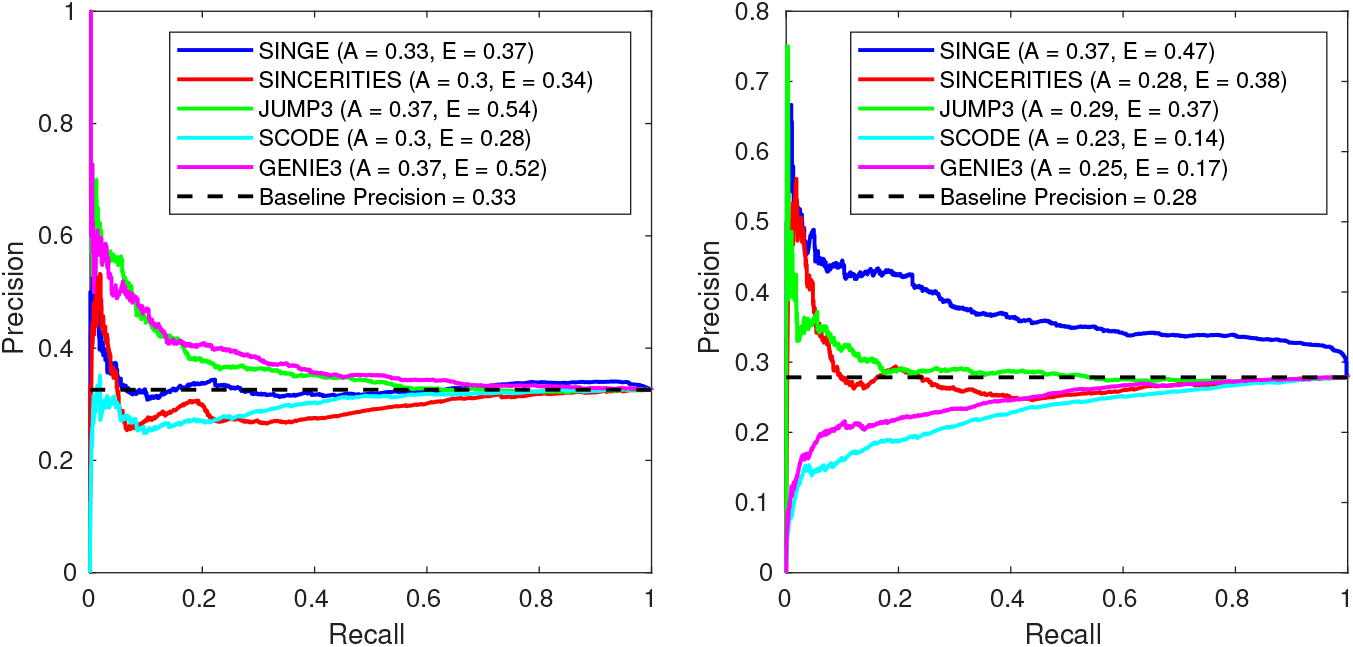
Precision-recall performance of network inference methods on the retinoic acid-driven differentiation dataset for ESCAPE gold standard interactions from (a) ChIP-chip/ChIP-seq and (b) *lof/gof* studies.

**Figure S4:**
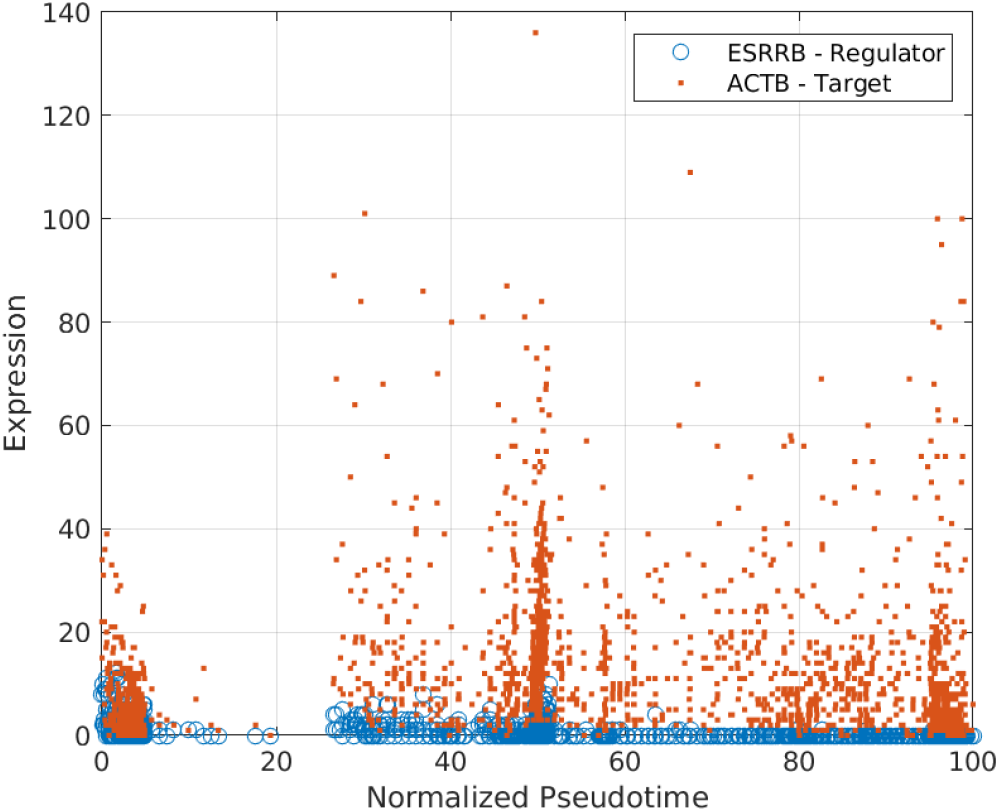
Expression trends of Esrrb and Actb show no apparent lag between regulator (Esrrb) and target expression (Actb). The interaction Esrrb→Actb is ranked highly by SCODE but not by SINGE.

**Figure S5:**
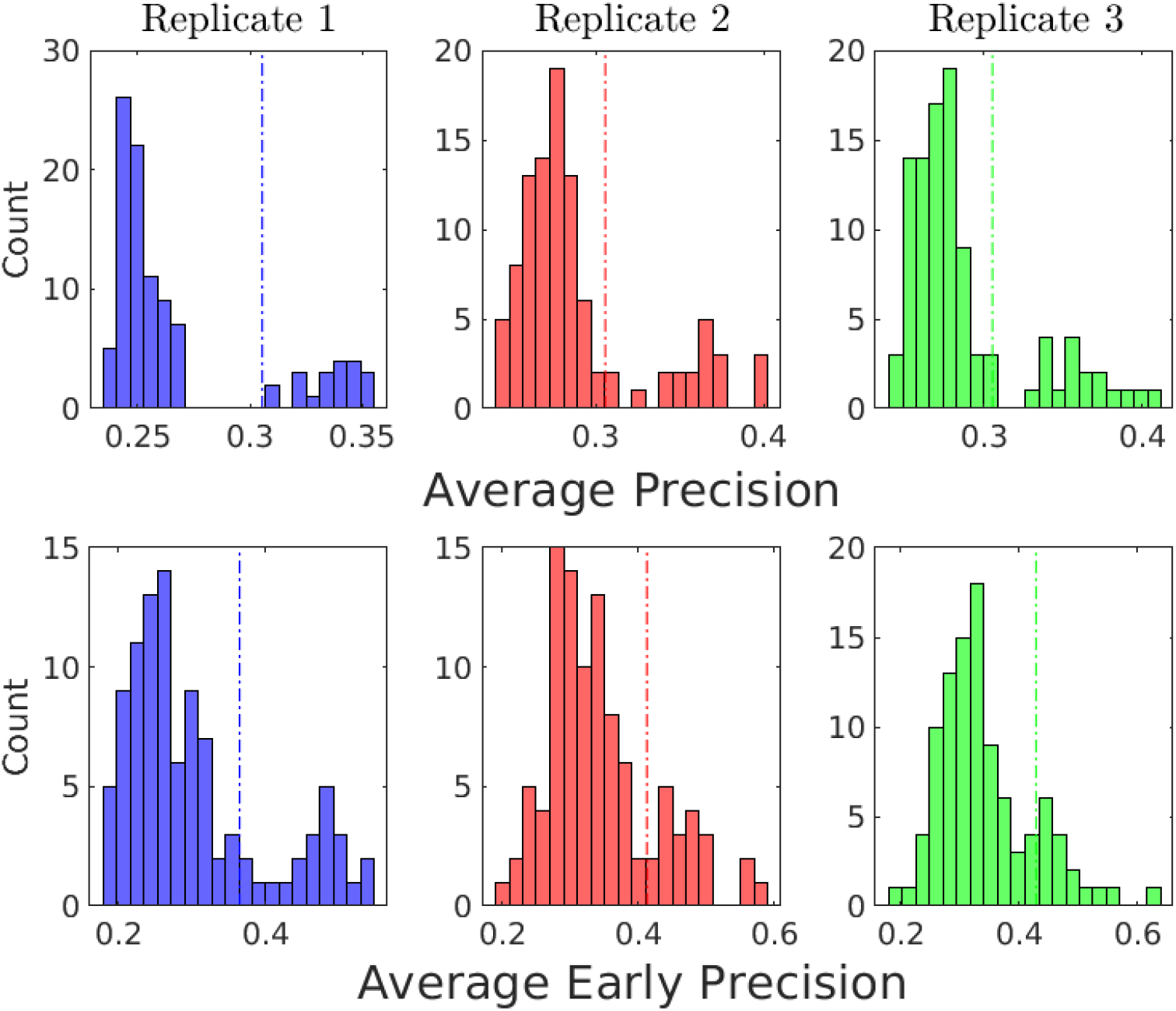
Histograms of average precision and average early precision for individual hyperparameters for the retinoic acid-driven differentiation dataset. The vertical lines depict the performance of the final SINGE GRN that ensembles all hyperparameter values.

**Figure S6:**
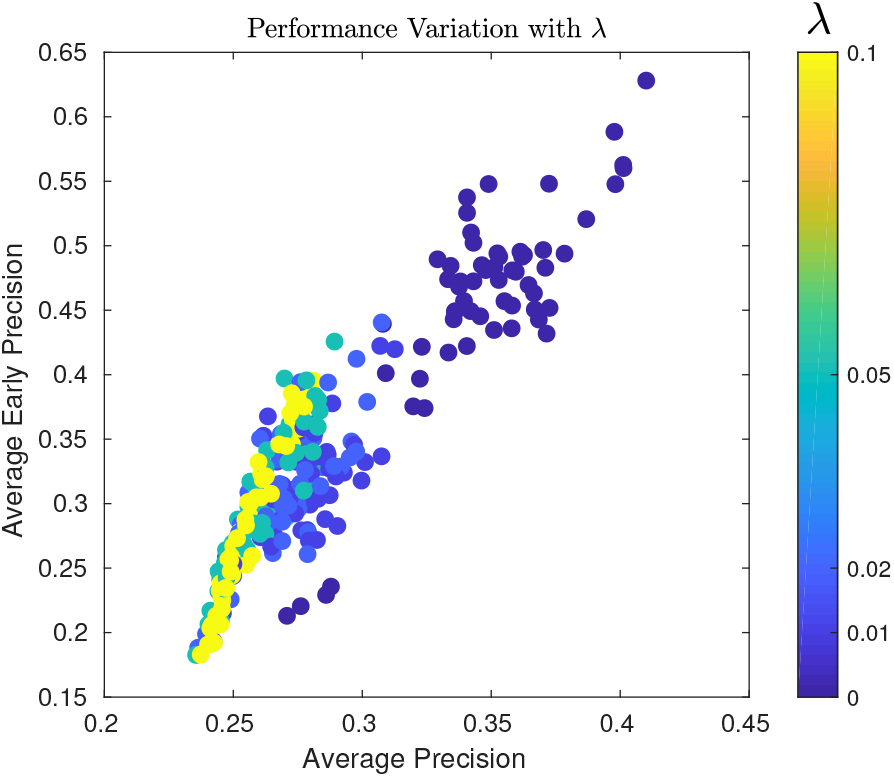
Scatter plot showing the effect of λ on the average precision and average early precision for the retinoic acid-driven differentiation dataset.

**Figure S7:**
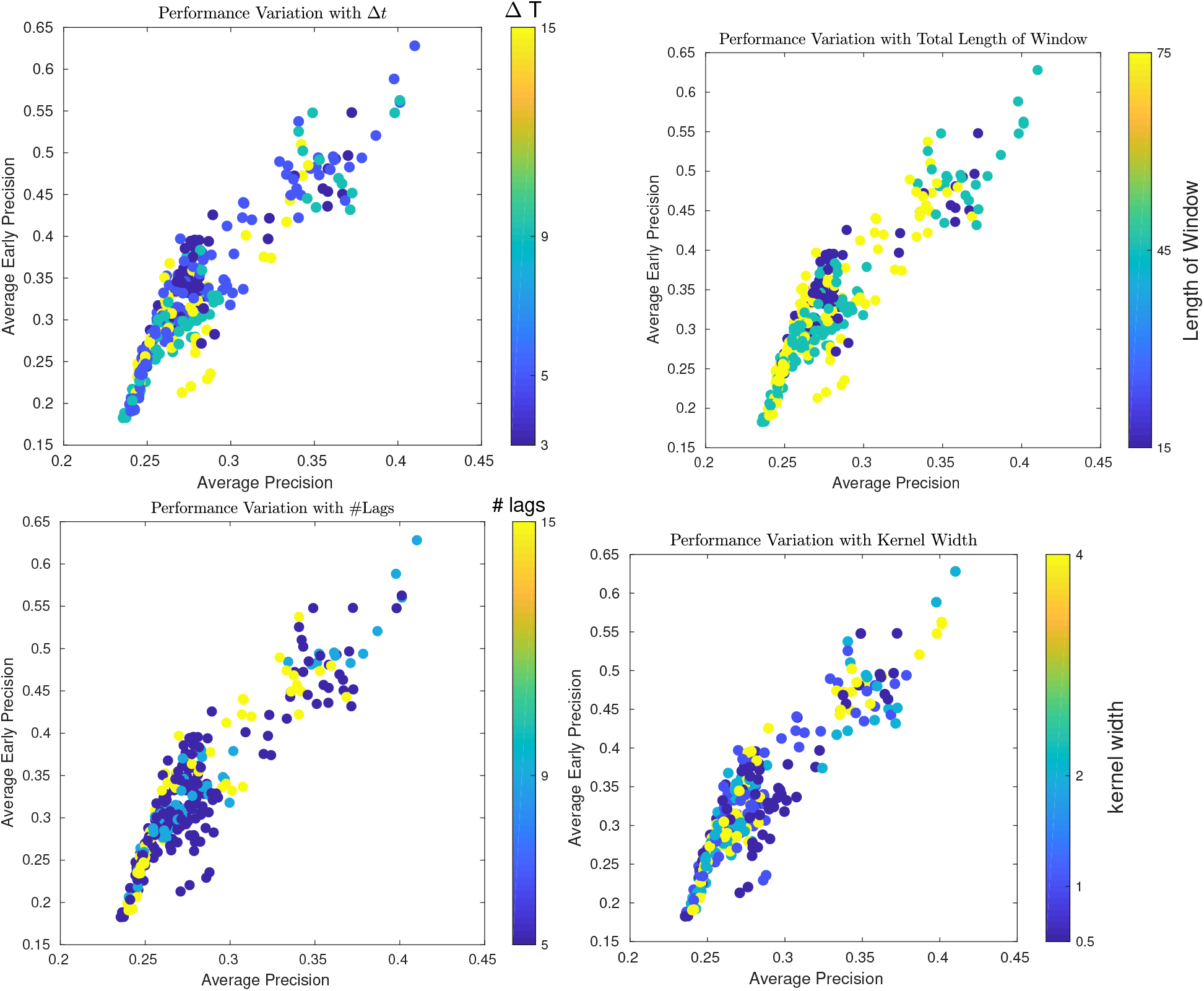
Scatter plot showing the effect of number of time resolution ΔT, the total length of the analysis window, the number of lagged coefficients, and the kernel width used in GLG on the average precision and average early precision for the retinoic acid-driven differentiation dataset.

**Figure S8:**
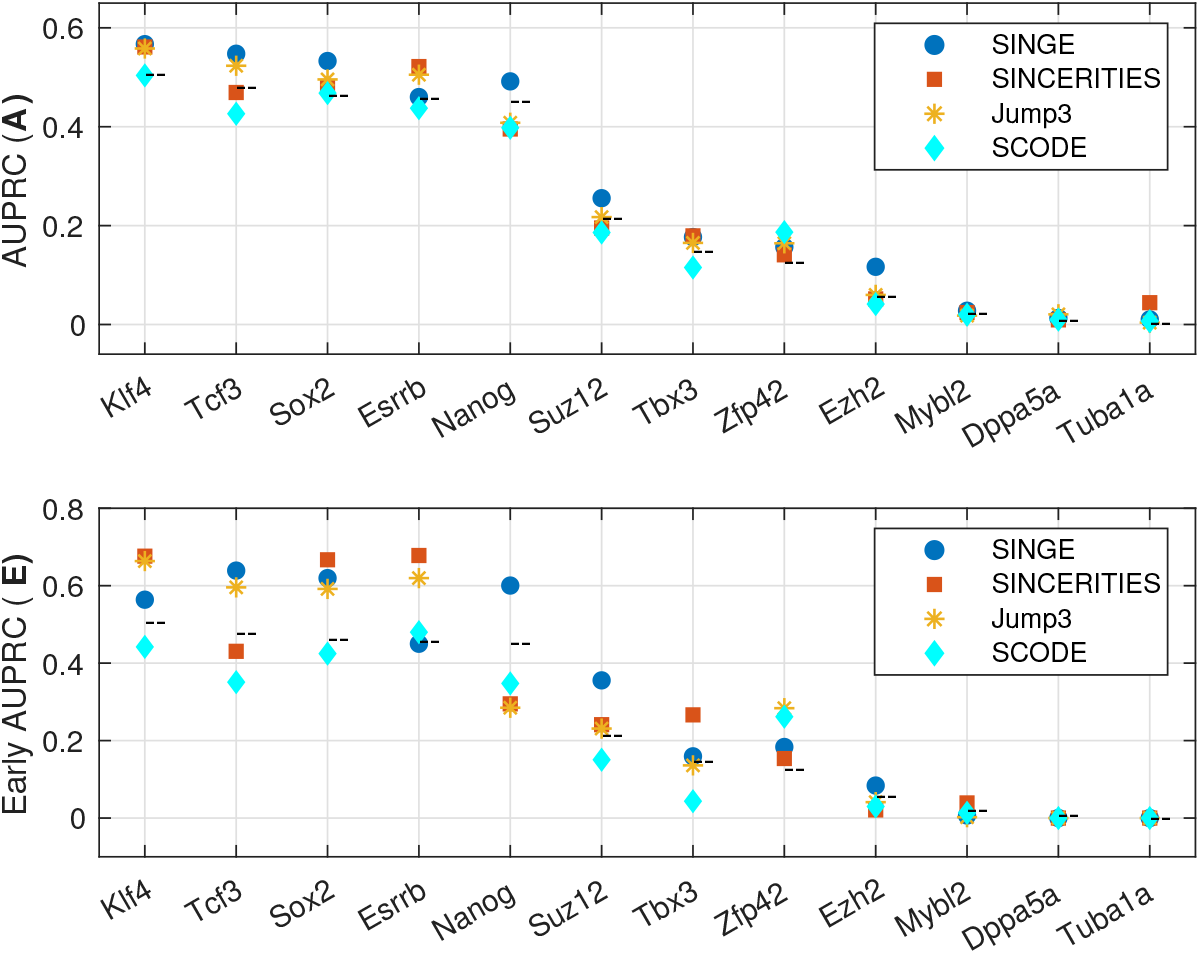
Average precision and average early precision evaluated for individual regulators for rankings obtained using the *Order Only* dataset obtained from Monocle 2. The dashed line (––) indicates random performance. The Jump3 results are the same as in Figure 6 because it does not use pseudotime values.

**Figure S9:**
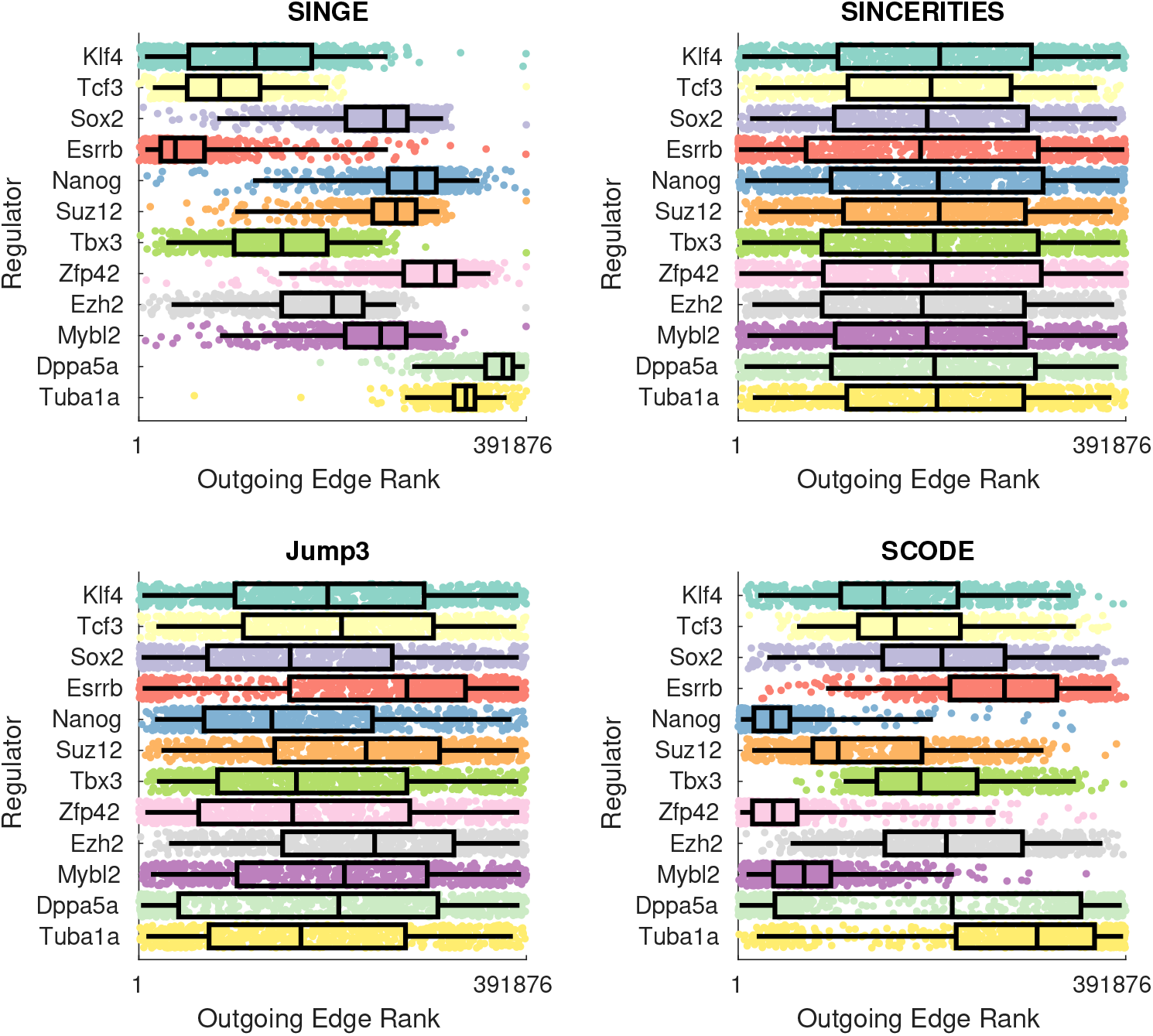
Boxplots of outgoing edge ranks for each regulator in each predicted GRN obtained using the *Order Only* dataset obtained from Monocle 2. The Jump3 results are the same as in Figure 7 because it does not use pseudotime values.

**Figure S10:**
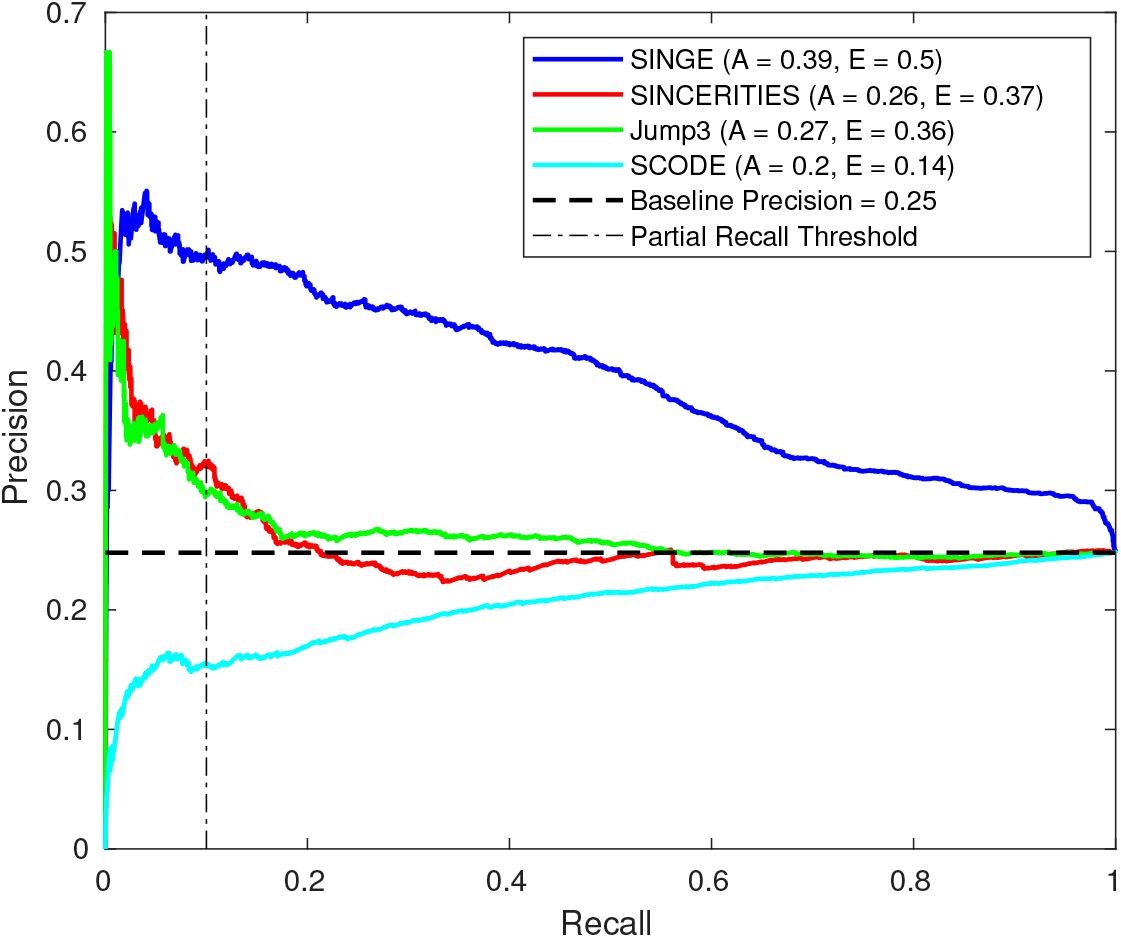
Precision-recall performance comparison of the four methods when using *Order Only* dataset obtained from Monocle 2. Key: **A** - Average Precision, **E** - Average Early Precision (≤ 0.1 recall). The Jump3 results are the same as in Figure 5 because it does not use pseudotime values.

**Figure S11:**
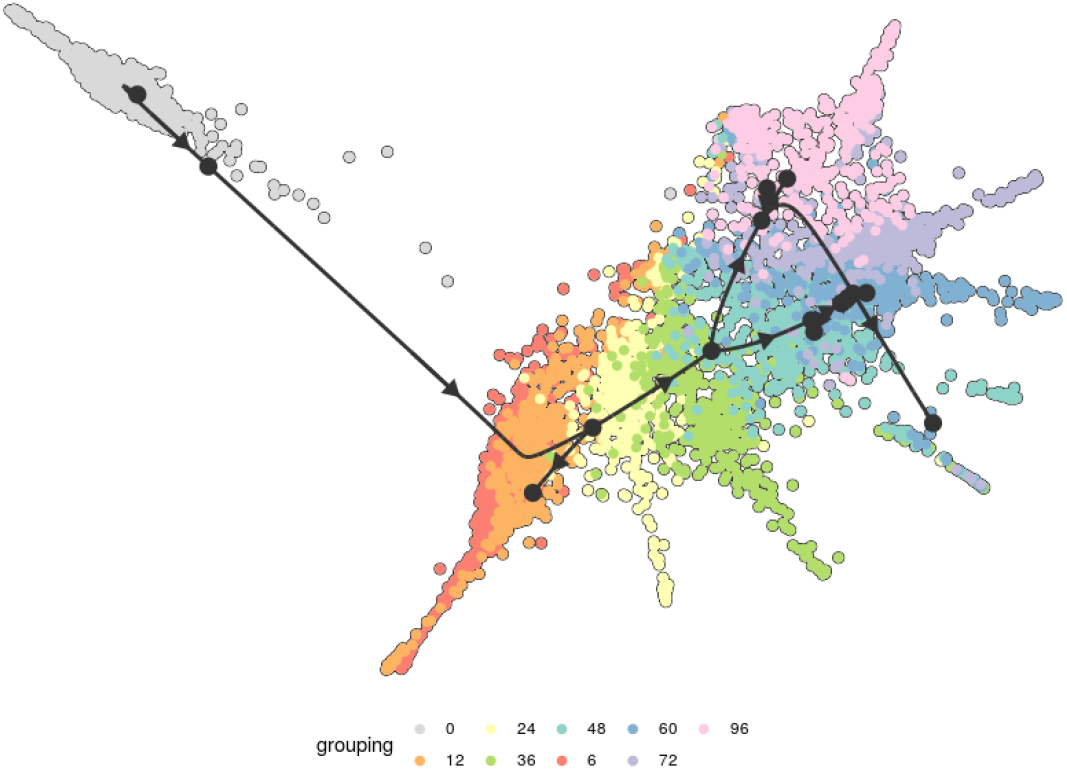
Trajectory inference using 626 genes from the retinoic acid-driven differentiation dataset using PAGA Tree. We select the longest branch, with 2631 cells (737 of which are common with the Monocle 2 branch studied) for our analysis.

**Figure S12:**
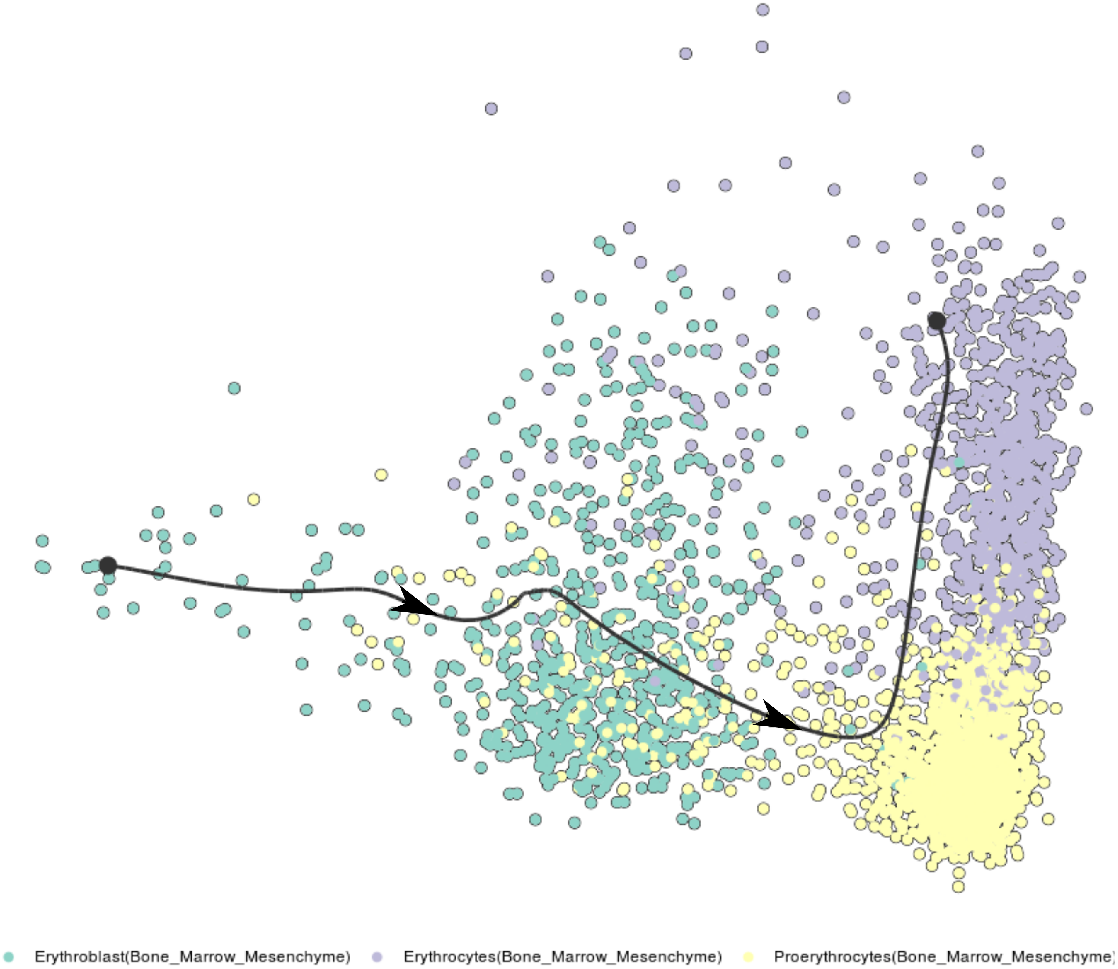
Trajectory inference from the mouse bone marrow dataset using Embeddr. The dataset has 3025 genes, 3105 cells, and a linear trajectory.

## Footnotes

1 The default recommendation for using SINGE with count-based data would be to use the hyperparameter ‘`--family poisson`.’ However, we discovered that the *glmnet* package for MATLAB suffers a high rate of memory segmentation violations when invoked for larger datasets with the Poisson distribution. Log-transforming the count-based data and using the hyperparameter ‘`--family gaussian`’ mitigates this issue.

## References

[1] A. Tanay and A. Regev, “Scaling single-cell genomics from phenomenology to mechanism,” Nature,vol. 541, no. 7637, p. 331, 2017.

[2] C. Trapnell, “Defining cell types and states with single-cell genomics,” Genome Research,vol. 25, no. 10, pp. 1491–1498, 2015.

[3] R. Bacher and C. Kendziorski, “Design and computational analysis of single-cell RNA-sequencing experiments,” Genome Biology,vol. 17, no. 1, p. 63, 2016.

[4] M. W. Fiers, L. Minnoye, S. Aibar, C. Bravo Gonzalez-Blas, Z. Kalender Atak, and S. Aerts, “Mapping gene regulatory networks from single-cell omics data,” Briefings in Functional Genomics, 2018.

[5] M. Blencowe, D. Arneson, J. Ding, Y.-W. Chen, Z. Saleem, and X. Yang, “Network modeling of single-cell omics data: challenges, opportunities, and progresses,” Emerging Topics in Life Sciences,vol. 3, pp. 379–398, Aug. 2019.

[6] H. Nguyen, D. Tran, B. Tran, B. Pehlivan, and T. Nguyen, “A comprehensive survey of regulatory network inference methods using single cell RNA sequencing data,” Briefings in Bioinformatics, Sept. 2020.

[7] R. De Smet and K. Marchal, “Advantages and limitations of current network inference methods,” Nature Reviews Microbiology,vol. 8, no. 10, p. 717, 2010.

[8] D. Marbach, J. C. Costello, R. Kuffner, N. M. Vega, R. J. Prill, D. M. Camacho, K. R. Allison, A. Aderhold, R. Bonneau, Y. Chen, et al., “Wisdom of crowds for robust gene network inference,” Nature Methods,vol. 9, pp. 796–804, Aug 2012.

[9] D. Chasman, A. F. Siahpirani, and S. Roy, “Network-based approaches for analysis of complex biological systems,” Current Opinion in Biotechnology, 2016.

[10] T. E. Chan, M. P. Stumpf, and A. C. Babtie, “Gene regulatory network inference from single-cell data using multivariate information measures,” Cell S’ystems,vol. 5, no. 3, pp. 251–267, 2017.

[11] J. Intosalmi, H. Mannerstrom, S. Hiltunen, and H. Lahdesmaki, “SCHiRM: Single cell hierarchical regression model to detect dependencies in read count data,” bioRxiv, 2018.

[12] V. A. Huynh-Thu, A. Irrthum, L. Wehenkel, and P. Geurts, “Inferring Regulatory Networks from Expression Data Using Tree-Based Methods,” PLoS ONE,vol. 5, p. e12776, sep 2010.

[13] Z. Bar-Joseph, A. Gitter, and I. Simon, “Studying and modelling dynamic biological processes using time-series gene expression data,” Nature Reviews Genetics,vol. 13, no. 8, p. 552, 2012.

[14] V. A. Huynh-Thu and G. Sanguinetti, “Combining tree-based and dynamical systems for the inference of gene regulatory networks,” Bioinformatics,vol. 31, no. 10, pp. 1614–1622, 2015.

[15] H. Matsumoto, H. Kiryu, C. Furusawa, M. S. Ko, S. B. Ko, N. Gouda, T. Hayashi, and I. Nikaido, “SCODE: An efficient regulatory network inference algorithm from single-cell RNA-seq during differentiation,” Bioinformatics,p. btx194, 2017.

[16] N. P. Gao, S. M. Ud-Dean, and R. Gunawan, “Gene regulatory network inference using time-stamped cross-sectional single cell expression data,” IFAC-PapersOnLine,vol. 49, no. 26, pp. 147–152, 2016.

[17] A. Gitter, “Single-cell RNA-seq pseudotime estimation algorithms,” doi:10.5281/zenodo.1297422, Jun 2018.

[18] W. Saelens, R. Cannoodt, H. Todorov, and Y. Saeys, “A comparison of single-cell trajectory inference methods,” Nature Biotechnology,vol. 37, pp. 547–554, May 2019.

[19] N. Leng, L.-F. Chu, C. Barry, Y. Li, J. Choi, X. Li, P. Jiang, R. M. Stewart, J. A. Thomson, and C. Kendziorski, “Oscope identifies oscillatory genes in unsynchronized single-cell RNA-seq experiments,” Nature Methods,vol. 12, no. 10, pp. 947–950, 2015.

[20] Z. Liu, H. Lou, K. Xie, H. Wang, N. Chen, O. M. Aparicio, M. Q. Zhang, R. Jiang, and T. Chen, “Reconstructing cell cycle pseudo time-series via single-cell transcriptome data,” Nature Communications,vol. 8, no. 1, p. 22, 2017.

[21] S. C. Bendall, K. L. Davis, E.-a. D. Amir, M. D. Tadmor, E. F. Simonds, T. J. Chen, D. K. Shenfeld, G. P. Nolan, and D. Peer, “Single-cell trajectory detection uncovers progression and regulatory coordination in human B cell development,” Cell, vol. 157, no. 3, pp. 714–725, 2014.

[22] J. Shin, D. Berg, Y. Zhu, J. Shin, J. Song, M. Bonaguidi, G. Enikolopov, D. Nauen, K. Christian, G.-l. Ming, and H. Song, “Single-cell RNA-Seq with Waterfall reveals molecular cascades underlying adult neurogenesis,” Cell Stem Cell,vol. 17, pp. 360–372, Sep 2015.

[23] M. Setty, M. D. Tadmor, S. Reich-Zeliger, O. Angel, T. M. Salame, P. Kathail, K. Choi, S. Bendall, N. Friedman, and D. Pe'er, “Wishbone identifies bifurcating developmental trajectories from single-cell data,” Nature Biotechnology,vol. 34, no. 6, pp. 637–645, 2016.

[24] H. Matsumoto and H. Kiryu, “SCOUP: a probabilistic model based on the Ornstein–Uhlenbeck process to analyze single-cell expression data during differentiation,” BMC Bioinformatics,vol. 17, no. 1, p. 232, 2016.

[25] X. Qiu, Q. Mao, Y. Tang, L. Wang, R. Chawla, H. A. Pliner, and C. Trapnell, “Reversed graph embedding resolves complex single-cell trajectories,” Nature Methods,vol. 14, no. 10, p. 979, 2017.

[26] J. Zhang, T. Zhou, and Q. Nie, “Topographer reveals dynamic mechanisms of cell fate decisions from single-cell transcriptomic data,” bioRxiv, 2018.

[27] N. Papili Gao, S. M. Ud-Dean, O. Gandrillon, and R. Gunawan, “Sincerities: inferring gene regulatory networks from time-stamped single cell transcriptional expression profiles,” Bioinformatics, vol. 34, no. 2, pp. 258–266, 2017.

[28] P.-C. Aubin-Frankowski and J.-P. Vert, “Gene regulation inference from single-cell RNA-seq data with linear differential equations and velocity inference,” bioRxiv, 2018.

[29] A. Ocone, L. Haghverdi, N. S. Mueller, and F. J. Theis, “Reconstructing gene regulatory dynamics from high-dimensional single-cell snapshot data,” Bioinformatics,vol. 31, no. 12, pp. i89–i96, 2015.

[30] J. Wei, X. Hu, X. Zou, and T. Tian, “Reverse-engineering of gene networks for regulating early blood development from single-cell measurements,” BMC Medical Genomics,vol. 10, no. 5, p. 72, 2017.

[31] A. T. Specht and J. Li, “Leap: constructing gene co-expression networks for single-cell RNA-sequencing data using pseudotime ordering,” Bioinformatics,vol. 33, no. 5, pp. 764–766, 2016.

[32] M. Sanchez-Castillo, D. Blanco, I. M. Tienda-Luna, M. Carrion, and Y. Huang, “A Bayesian framework for the inference of gene regulatory networks from time and pseudo-time series data,” Bioinformatics,vol. 34, no. 6, pp. 964–970, 2017.

[33] X. Qiu, A. Rahimzamani, L. Wang, Q. Mao, T. Durham, J. L. McFaline-Figueroa, L. Saunders, C. Trapnell, and S. Kannan, “Towards inferring causal gene regulatory networks from single cell expression measurements,” bioRxiv, 2018.

[34] P. Tsakanikas, D. V. Manatakis, and E. S. Manolakos, “Machine learning methods to reverse engineer dynamic gene regulatory networks governing cell state transitions,” bioRxiv, 2018.

[35] T. E. Chan, A. Pallaseni, A. C. Babtie, K. McEwen, and M. P. Stumpf, “Empirical Bayes meets information theoretical network reconstruction from single cell data,” bioRxiv, 2018.

[36] A. Bonnaffoux, U. Herbach, A. Richard, A. Guillemin, S. Giraud, P.-A. Gros, and O. Gandrillon, “Wasabi: a dynamic iterative framework for gene regulatory network inference,” bioRxiv, 2018.

[37] J. Kim, S. T. Jakobsen, K. N. Natarajan, and K. J. Won, “Gene network reconstruction using single cell transcriptomic data reveals key factors for embryonic stem cell differentiation,” bioRxiv, p. 2019.12.20.884163, Dec. 2019.

[38] P. Cordero and J. M. Stuart, “Tracing co-regulatory network dynamics in noisy, single-cell transcriptome trajectories,” in Pacific Symposium on Biocomputing 2017,pp. 576–587, World Scientific, 2017.

[39] C. W. Granger, “Investigating causal relations by econometric models and cross-spectral methods,” Econometrica: Journal of the Econometric Society,pp. 424–438, 1969.

[40] C. W. J. Granger, “Testing for causality: A personal viewpoint,” Journal of Economic Dynamics and Control,vol. 2, pp. 329–352, Jan. 1980.

[41] A. Fujita, P. Severino, J. R. Sato, and S. Miyano, “Granger causality in systems biology: Modeling gene networks in time series microarray data using vector autoregressive models,” in Brazilian Symposium on Bioinformatics,pp. 13–24, Springer, 2010.

[42] N. D. Mukhopadhyay and S. Chatterjee, “Causality and pathway search in microarray time series experiment,” Bioinformatics,vol. 23, no. 4, pp. 442–449, 2006.

[43] A. Shojaie and G. Michailidis, “Discovering graphical granger causality using the truncating lasso penalty,” Bioinformatics,vol. 26, no. 18, pp. i517–i523, 2010.

[44] J. D. Finkle, J. J. Wu, and N. Bagheri, “Windowed Granger causal inference strategy improves discovery of gene regulatory networks,” Proceedings of the National Academy of Sciences,vol. 115, no. 9, pp. 2252–2257, 2018.

[45] S. Heerah, R. Molinari, S. Guerrier, and A. Marshall-Colon, “Granger-Causal Testing for Irregularly Sampled Time Series with Application to Nitrogen Signaling in Arabidopsis,” bioRxiv, p. 2020.06.15.152819, June 2020.

[46] J. Lu, B. Dumitrascu, I. C. McDowell, B. Jo, A. Barrera, L. K. Hong, S. M. Leichter, T. E. Reddy, and B. E. Engelhardt, “Causal network inference from gene transcriptional time-series response to glucocorticoids,” PLOS Computational Biology, vol. 17, p. e1008223, Jan. 2021.

[47] M. T. Bahadori and Y. Liu, “Granger causality analysis in irregular time series,” in Proceedings of the 2012 SIAM International Conference on Data Mining,pp. 660–671, 2012.

[48] A. Arnold, Y. Liu, and N. Abe, “Temporal causal modeling with graphical Granger methods,” in Proceedings of the 13th ACM SIGKDD International Conference on Knowledge Discovery and Data mining,pp. 66–75, ACM, 2007.

[49] D. T. Gillespie, “Exact stochastic simulation of coupled chemical reactions,” The Journal of Physical Chemistry,vol. 81, pp. 2340–2361, Dec. 1977.

[50] M. Thattai and A. v. Oudenaarden, “Intrinsic noise in gene regulatory networks,” Proceedings of the National Academy of Sciences,vol. 98, pp. 8614–8619, July 2001.

[51] T. Hayashi, H. Ozaki, Y. Sasagawa, M. Umeda, H. Danno, and I. Nikaido, “Single-cell full-length total RNA sequencing uncovers dynamics of recursive splicing and enhancer RNAs,” Nature Communications, vol. 9, no. 1, p. 619, 2018.

[52] H. Xu, C. Baroukh, R. Dannenfelser, E. Y. Chen, C. M. Tan, Y. Kou, Y. E. Kim, I. R. Lemischka, and A. Ma'ayan, “Escape: database for integrating high-content published data collected from human and mouse embryonic stem cells,” Database, vol. 2013, 2013.

[53] DREAM4 In Silico Network Challenge. http://dreamchallenges.org/project/dream4-in-silico-network-challenge/.

[54] S. Semrau, J. E. Goldmann, M. Soumillon, T. S. Mikkelsen, R. Jaenisch, and A. Van Oudenaarden, “Dynamics of lineage commitment revealed by single-cell transcriptomics of differentiating embryonic stem cells,” Nature Communications,vol. 8, no. 1, p. 1096, 2017.

[55] J. Reimand, T. Arak, P. Adler, L. Kolberg, S. Reisberg, H. Peterson, and J. Vilo, “g:Profiler-a web server for functional interpretation of gene lists (2016 update),” Nucleic Acids Research,vol. 44, no. W1, pp. W83–W89, 2016.

[56] S. M. Morris, M. D. Tallquist, C. O. Rock, and J. A. Cooper, “Dual roles for the Dab2 adaptor protein in embryonic development and kidney transport,” The EMBO Journal,vol. 21, pp. 1555–1564, Apr 2002.

[57] B. Feldman, W. Poueymirou, V. E. Papaioannou, T. M. DeChiara, and M. Goldfarb, “Requirement of FGF-4 for postimplantation mouse development,” Science, vol. 267, no. 5195, pp. 246–249, 1995.

[58] I. Leaf, J. Tennessen, M. Mukhopadhyay, H. Westphal, and W. Shawlot, “Sfrp5 is not essential for axis formation in the mouse,” Genesis,vol. 44, pp. 573–578, Dec 2006.

[59] C. Meno, K. Gritsman, S. Ohishi, Y. Ohfuji, E. Heckscher, K. Mochida, A. Shimono, H. Kondoh, W. S. Talbot, E. J. Robertson, A. F. Schier, and H. Hamada, “Mouse Lefty2 and Zebrafish Antivin are feedback inhibitors of nodal signaling during vertebrate gastrulation,” Molecular Cell,vol. 4, pp. 287–298, Sep 1999.

[60] J. R. Barrow and M. R. Capecchi, “Targeted disruption of the Hoxb-2 locus in mice interferes with expression of Hoxb-1 and Hoxb-4,” Development,vol. 122, no. 12, pp. 3817–3828, 1996.

[61] E. E. Morrisey, Z. Tang, K. Sigrist, M. M. Lu, F. Jiang, H. S. Ip, and M. S. Parmacek, “GATA6 regulates HNF4 and is required for differentiation of visceral endoderm in the mouse embryo,” Genes & Development,vol. 12, pp. 3579–3590, Nov 1998.

[62] G. L. Radice, H. Rayburn, H. Matsunami, K. A. Knudsen, M. Takeichi, and R. O. Hynes, “Developmental defects in mouse embryos lacking N-Cadherin,” Developmental Biology,vol. 181, pp. 64–78, Jan 1997.

[63] W. C. Skarnes, B. Rosen, A. P. West, M. Koutsourakis, W. Bushell, V. Iyer, A. O. Mujica, M. Thomas, J. Harrow, T. Cox, D. Jackson, J. Severin, P. Biggs, J. Fu, M. Nefedov, P. J. de Jong, A. F. Stewart, and A. Bradley, “A conditional knockout resource for the genome-wide study of mouse gene function,” Nature,vol. 474, pp. 337–342, Jun 2011.

[64] J. Parant, A. Chavez-Reyes, N. A. Little, W. Yan, V. Reinke, A. G. Jochemsen, and G. Lozano, “Rescue of embryonic lethality in Mdm4-null mice by loss of Trp53 suggests a nonoverlapping pathway with MDM2 to regulate p53,” Nature Genetics,vol. 29, pp. 92–95, Sep 2001.

[65] G. E. Olson, J. C. Whitin, K. E. Hill, V. P. Winfrey, A. K. Motley, L. M. Austin, J. Deal, H. J. Cohen, and R. F. Burk, “Extracellular glutathione peroxidase (Gpx3) binds specifically to basement membranes of mouse renal cortex tubule cells,” American Journal of Physiology-Renal Physiology,vol. 298, pp. F1244–F1253, May 2010.

[66] T. M. DeChiara, A. Efstratiadis, and E. J. Robertsen, “A growth-deficiency phenotype in heterozygous mice carrying an insulin-like growth factor II gene disrupted by targeting,” Nature,vol. 345, pp. 78–80, May 1990.

[67] P. Sicinski, J. L. Donaher, Y. Geng, S. B. Parker, H. Gardner, M. Y. Park, R. L. Robker, J. S. Richards, L. K. McGinnis, J. D. Biggers, J. J. Eppig, R. T. Bronson, S. J. Elledge, and R. A. Weinberg, “Cyclin D2 is an FSH-responsive gene involved in gonadal cell proliferation and oncogenesis,” Nature,vol. 384, pp. 470–474, Dec 1996.

[68] Y. Xiao, H. Ma, P. Wan, D. Qin, X. Wang, X. Zhang, Y. Xiang, W. Liu, J. Chen, Z. Yi, and L. Li, “Trp-Asp (WD) repeat domain 1 is essential for mouse peri-implantation development and regulates Cofilin phosphorylation,” The Journal of Biological Chemistry,vol. 292, pp. 1438–1448, Jan 2017.

[69] T. Sakai, S. Li, D. Docheva, C. Grashoff, K. Sakai, G. Kostka, A. Braun, A. Pfeifer, P. D. Yurchenco, and R. Fässler, “Integrin-linked kinase (ILK) is required for polarizing the epiblast, cell adhesion, and controlling actin accumulation,” Genes & Development,vol. 17, pp. 926–40, Apr 2003.

[70] J. Egea, C. Erlacher, E. Montanez, I. Burtscher, S. Yamagishi, M. Hess, F. Hampel, R. Sanchez, M. T. Rodriguez-Manzaneque, M. R. Böosl, R. Fässler, H. Lickert, and R. Klein, “Genetic ablation of FLRT3 reveals a novel morphogenetic function for the anterior visceral endoderm in suppressing mesoderm differentiation,” Genes & Development,vol. 22, pp. 3349–62, Dec 2008.

[71] V. E. Sollars, B. J. McEntee, J. B. Engiles, J. L. Rothstein, and A. M. Buchberg, “A novel transgenic line of mice exhibiting autosomal recessive male-specific lethality and non-alcoholic fatty liver disease,” Human Molecular Genetics,vol. 11, pp. 2777–2786, Oct 2002.

[72] A. C. Carpenter, S. Rao, J. M. Wells, K. Campbell, and R. A. Lang, “Generation of mice with a conditional null allele for Wntless,” Genesis,vol. 48, pp. 554–558, Aug 2010.

[73] Y. Wang, N. Thekdi, P. M. Smallwood, J. P. Macke, and J. Nathans, “Frizzled-3 is required for the development of major fiber tracts in the rostral CNS,” Journal of Neuroscience,vol. 22, pp. 8563–73, Oct 2002.

[74] P. Gorry, T. Lufkin, A. Dierich, C. Rochette-Egly, D. Décimo, P. Dolle, M. Mark, B. Durand, and P. Chambon, “The cellular retinoic acid binding protein I is dispensable,” Proceedings of the National Academy of Sciences of the United States of America,vol. 91, pp. 9032–6, Sep 1994.

[75] F. Kuusisto, J. Steill, Z. Kuang, J. Thomson, D. Page, and R. Stewart, “A simple text mining approach for ranking pairwise associations in biomedical applications,” AMIA Joint Summits on Translational Science proceedings. AMIA Joint Summits on Translational Science, vol. 2017, pp. 166–174, 2017.

[76] T. Kunath, M. K. Saba-El-Leil, M. Almousailleakh, J. Wray, S. Meloche, and A. Smith, “FGF stimulation of the Erk1/2 signalling cascade triggers transition of pluripotent embryonic stem cells from self-renewal to lineage commitment,” Development,vol. 134, no. 16, pp. 2895–2902, 2007.

[77] Y. Yamanaka, F. Lanner, and J. Rossant, “FGF signal-dependent segregation of primitive endoderm and epiblast in the mouse blastocyst,” Development,vol. 137, no. 5, pp. 715–724, 2010.

[78] D. Krawchuk, N. Honma-Yamanaka, S. Anani, and Y. Yamanaka, “FGF4 is a limiting factor controlling the proportions of primitive endoderm and epiblast in the ICM of the mouse blastocyst,” Developmental Biology,vol. 384, no. 1, pp. 65–71, 2013.

[79] X. Zhang, A. Friedman, S. Heaney, P. Purcell, and R. L. Maas, “Meis homeoproteins directly regulate Pax6 during vertebrate lens morphogenesis,” Genes & Development,vol. 16, no. 16, pp. 2097–2107, 2002.

[80] M. T. Pankratz, X.-J. Li, T. M. LaVaute, E. A. Lyons, X. Chen, and S.-C. Zhang, “Directed neural differentiation of human embryonic stem cells via an obligated primitive anterior stage,” Stem Cells,vol. 25, no. 6, pp. 1511–1520, 2007.

[81] D. Shimosato, M. Shiki, and H. Niwa, “Extra-embryonic endoderm cells derived from ES cells induced by GATA factors acquire the character of XEN cells,” BMC Developmental Biology,vol. 7, no. 1, p. 80, 2007.

[82] M. P. Stavridis, J. S. Lunn, B. J. Collins, and K. G. Storey, “A discrete period of FGF-induced Erk1/2 signalling is required for vertebrate neural specification,” Development,vol. 134, no. 16, pp. 2889–2894, 2007.

[83] K. R. Finley, J. Tennessen, and W. Shawlot, “The mouse secreted frizzled-related protein 5 gene is expressed in the anterior visceral endoderm and foregut endoderm during early post-implantation development,” Gene Expression Patterns,vol. 3, no. 5, pp. 681–684, 2003.

[84] K. Takaoka, H. Nishimura, and H. Hamada, “Both nodal signalling and stochasticity select for prospective distal visceral endoderm in mouse embryos,” Nature Communications,vol. 8, no. 1, p. 1492, 2017.

[85] K. Q. Cai, C. D. Capo-Chichi, M. E. Rula, D.-H. Yang, and X.-X. Xu, “Dynamic GATA6 expression in primitive endoderm formation and maturation in early mouse embryogenesis,” Developmental Dynamics,vol. 237, no. 10, pp. 2820–2829, 2008.

[86] J. Artus, P. Douvaras, A. Piliszek, J. Isern, M. H. Baron, and A.-K. Hadjantonakis, “BMP4 signaling directs primitive endoderm-derived XEN cells to an extraembryonic visceral endoderm identity,” Developmental Biology,vol. 361, no. 2, pp. 245–262, 2012.

[87] X. Han, R. Wang, Y. Zhou, L. Fei, H. Sun, S. Lai, A. Saadatpour, Z. Zhou, H. Chen, F. Ye, D. Huang, Y. Xu, W. Huang, M. Jiang, X. Jiang, J. Mao, Y. Chen, C. Lu, J. Xie, Q. Fang, Y. Wang, R. Yue, T. Li, H. Huang, S. H. Orkin, G.-C. Yuan, M. Chen, and G. Guo, “Mapping the mouse cell atlas by microwell-seq,” Cell, vol. 172, no. 5, pp. 1091–1107.e17, 2018.

[88] K. Campbell, C. P. Ponting, and C. Webber, “Laplacian eigenmaps and principal curves for high resolution pseudotemporal ordering of single-cell rna-seq profiles,” bioRxiv, 2015.

[89] J. Li, F. He, P. Zhang, S. Chen, H. Shi, Y. Sun, G. Ying, H. Yang, S. D. Nimer, Q.-F. Wang, M. Xu, and F.-C. Yang, “ASXL2 Is Required for Normal Hematopoiesis and Loss of asxl2 Leads to Myeloid Malignancies in Mice,” Blood,vol. 128, pp. 1509–1509, 12 2016.

[90] J.-B. Micol, A. Pastore, D. Inoue, N. Duployez, E. Kim, S. C.-W. Lee, B. H. Durham, Y. R. Chung, H. Cho, X. J. Zhang, A. Yoshimi, A. Krivtsov, R. Koche, E. Solary, A. Sinha, C. Preudhomme, and O. Abdel-Wahab, “Asxl2 is essential for haematopoiesis and acts as a haploinsufficient tumour suppressor in leukemia,” Nature Communications,vol. 8, p. 15429, 2017.

[91] J. C. W. Marsh, F. Gutierrez-Rodrigues, J. Cooper, J. Jiang, S. Gandhi, S. Kajigaya, X. Feng, M. d. P. F. Ibanez, F. S. Donaires, J. P. Lopes da Silva, Z. Li, S. Das, M. Ibanez, A. E. Smith, N. Lea, S. Best, R. Ireland, A. G. Kulasekararaj, D. P. McLornan, A. Pagliuca, I. Callebaut, N. S. Young, R. T. Calado, D. M. Townsley, and G. J. Mufti, “Heterozygous RTEL1 variants in bone marrow failure and myeloid neoplasms,” Blood Advances,vol. 2, pp. 36–48, Jan 2018.

[92] A. Balakumaran, P. J. Mishra, E. Pawelczyk, S. Yoshizawa, B. J. Sworder, N. Cherman, S. A. Kuznetsov, P. Bianco, N. Giri, S. A. Savage, G. Merlino, B. Dumitriu, C. E. Dunbar, N. S. Young, B. P. Alter, and P. G. Robey, “Bone marrow skeletal stem/progenitor cell defects in dyskeratosis congenita and telomere biology disorders,” Blood,vol. 125, pp. 793–802, Jan 2015.

[93] R. Cannoodt, W. Saelens, L. Deconinck, and Y. Saeys, “dyngen: a multi-modal simulator for spearheading new single-cell omics analyses,” bioRxiv,p. 2020.02.06.936971, Feb. 2020.

[94] D. Risso, J. Ngai, T. P. Speed, and S. Dudoit, “Normalization of RNA-seq data using factor analysis of control genes or samples,” Nature Biotechnology,vol. 32, no. 9, pp. 896–902, 2014.

[95] F. A. Wolf, F. K. Hamey, M. Plass, J. Solana, J. S. Dahlin, B. Gättgens, N. Rajewsky, L. Simon, and F. J. Theis, “Paga: graph abstraction reconciles clustering with trajectory inference through a topology preserving map of single cells,” Genome Biology,vol. 20, no. 1, p. 59, 2019.

[96] R. Pordes, D. Petravick, B. Kramer, D. Olson, M. Livny, A. Roy, P. Avery, K. Blackburn, T. Wenaus, F. Wuärthwein, I. Foster, R. Gardner, M. Wilde, A. Blatecky, J. McGee, and R. Quick, “The open science grid,” Journal of Physics: Conference Series,vol. 78, p. 012057, Jul 2007.

[97] R. A. Erickson, M. N. Fienen, S. G. McCalla, E. L. Weiser, M. L. Bower, J. M. Knudson, and G. Thain, “Wrangling distributed computing for high-throughput environmental science: An introduction to HTCondor,” PLoS Computational Biology,vol. 14, no. 10, p. e1006468, 2018.

[98] H. Hu, Y.-R. Miao, L.-H. Jia, Q.-Y. Yu, Q. Zhang, and A.-Y. Guo, “AnimalTFDB 3.0: a comprehensive resource for annotation and prediction of animal transcription factors,” Nucleic Acids Research, vol. 47, pp. D33–D38, 09 2018.

[99] J. Qian, T. Hastie, J. Friedman, R. Tibshirani, and N. Simon, “GLMNET for MATLAB.” http://www.stanford.edu/~hastie/glmnet_matlab/, 2013.

[100] G. C. Linderman, J. Zhao, and Y. Kluger, “Zero-preserving imputation of scRNA-seq data using low-rank approximation,” bioRxiv, 2018.

[101] D. van Dijk, R. Sharma, J. Nainys, K. Yim, P. Kathail, A. J. Carr, C. Burdziak, K. R. Moon, C. L. Chaffer, D. Pattabiraman, B. Bierie, L. Mazutis, G. Wolf, S. Krishnaswamy, and D. Pe’er, “Recovering Gene Interactions from Single-Cell Data Using Data Diffusion,” Cell,vol. 174, pp. 716–729.e27, July 2018.

[102] T. Andrews and M. Hemberg, “False signals induced by single-cell imputation [version 1; referees: 4 approved with reservations],” F1000Research,vol. 7, no. 1740, 2018.

[103] L. Zhang and S. Zhang, “Comparison of computational methods for imputing single-cell RNA-sequencing data,” IEEE/ACM Transactions on Computational Biology and Bioinformatics, pp. 376–389, 2018.

[104] L. Breiman, “Bagging predictors,” Machine Learning,vol. 24, pp. 123–140, Aug 1996.

[105] M. van Erp and L. Schomaker, “Variants of the Borda count method for combining ranked classifier hypotheses,” in Proceedings 7th International Workshop on Frontiers in Handwriting Recognition (7th IWFHR) (L. Schomaker and L. Vuurpijl, eds.), pp. 443–452, International Unipen Foundation, 2000.

[106] J. Fraenkel and B. Grofman, “The Borda Count and its real-world alternatives: Comparing scoring rules in Nauru and Slovenia,” Australian Journal of Political Science,vol. 49, pp. 186–205, Apr. 2014.

[107] N. Meinshausen and P. Buhlmann, “Stability selection,” Journal of the Royal Statistical Society: Series B (Statistical Methodology),vol. 72, no. 4, pp. 417–473, 2010.

[108] A.-C. Haury, F. Mordelet, P. Vera-Licona, and J.-P. Vert, “TIGRESS: Trustful Inference of Gene REgulation using Stability Selection,” BMC Systems Biology,vol. 6, p. 145, Nov. 2012.

[109] M. E. Ahsen, R. Vogel, and G. Stolovitzky, “Unsupervised evaluation and weighted aggregation of ranked predictions,” arXiv, Feb 2018.

[110] K. Street, D. Risso, R. B. Fletcher, D. Das, J. Ngai, N. Yosef, E. Purdom, and S. Dudoit, “Slingshot: cell lineage and pseudotime inference for single-cell transcriptomics,” BMC Genomics, vol. 19, no. 1, p. 477, 2018.

[111] R. Cannoodt, W. Saelens, D. Sichien, S. Tavernier, S. Janssens, M. Guilliams, B. Lambrecht, K. D. Preter, and Y. Saeys, “Scorpius improves trajectory inference and identifies novel modules in dendritic cell development,” bioRxiv, 2016.

[112] M. D. Luecken and F. J. Theis, “Current best practices in single-cell RNA-seq analysis: a tutorial,” Molecular Systems Biology,vol. 15, no. 6, p. e8746, 2019.

[113] S. Aibar, C. B. Gonzalez-Blas, T. Moerman, V. A. Huynh-Thu, H. Imrichova, G. Hulselmans, F. Rambow, J.-C. Marine, P. Geurts, J. Aerts, J. van den Oord, Z. K. Atak, J. Wouters, and S. Aerts, “SCENIC: single-cell regulatory network inference and clustering,” Nature Methods,vol. 14, pp. 1083–1086, nov 2017.

[114] M. Ashburner, C. A. Ball, J. A. Blake, D. Botstein, H. Butler, J. M. Cherry, A. P. Davis, K. Dolinski, S. S. Dwight, J. T. Eppig, M. A. Harris, D. P. Hill, L. Issel-Tarver, A. Kasarskis, S. Lewis, J. C. Matese, J. E. Richardson, M. Ringwald, G. M. Rubin, and G. Sherlock, “Gene Ontology: tool for the unification of biology. The Gene Ontology Consortium,” Nature Genetics, vol. 25, pp. 25–29, May 2000.

[115] C. J. Bult, J. A. Blake, C. L. Smith, J. A. Kadin, J. E. Richardson, and the Mouse Genome Database Group, “Mouse Genome Database (MGD) 2019,” Nucleic Acids Research,vol. 47, no. D1, pp. D801–D806, 2019.

[116] M. T. Bahadori and Y. Liu, “An examination of practical Granger causality inference,” in Proceedings of the 2013 SIAM International Conference on Data Mining,pp. 467–475, 2013.

[117] P. A. Valdes-Sosa, J. M. Sanchez-Bornot, A. Lage-Castellanos, M. Vega-Hernandez, J. Bosch-Bayard, L. Melie-García, and E. Canales-Rodríguez, “Estimating brain functional connectivity with sparse multivariate autoregression,” Philosophical Transactions of the Royal Society of London B: Biological Sciences,vol. 360, no. 1457, pp. 969–981, 2005.

[118] J. Swift and G. M. Coruzzi, “A matter of time — How transient transcription factor interactions create dynamic gene regulatory networks,” Biochimica et Biophysica Acta (BBA) - Gene Regulatory Mechanisms,vol. 1860, pp. 75–83, Jan. 2017.

[119] A. F. Siahpirani and S. Roy, “A prior-based integrative framework for functional transcriptional regulatory network inference,” Nucleic Acids Research,vol. 45, no. 4, pp. e21–e21, 2016.

[120] A. Greenfield, C. Hafemeister, and R. Bonneau, “Robust data-driven incorporation of prior knowledge into the inference of dynamic regulatory networks,” Bioinformatics,vol. 29, no. 8, pp. 1060–1067, 2013.

[121] C. Jansen, R. Ramirez, N. El-Ali, D. Gomez-Cabrero, J. Tegner, M. Merkenschlager, A. Conesa, and A. Mortazavi, “Building gene regulatory networks from single-cell ATAC-seq and RNA-seq using linked self-organizing maps,” bioRxiv, 2018.

[122] C. Burdziak, E. Azizi, S. Prabhakaran, and D. Pe'er, “A Nonparametric Multi-view Model for Estimating Cell Type-Specific Gene Regulatory Networks,” arXiv:1902.08138 [cs, q-bio, stat], Feb. 2019.

[123] J. Ding, B. J. Aronow, N. Kaminski, J. Kitzmiller, J. A. Whitsett, and Z. Bar-Joseph, “Reconstructing differentiation networks and their regulation from time series single-cell expression data,” Genome Research, 2018.

[124] M. Ciofani, A. Madar, C. Galan, M. Sellars, K. Mace, F. Pauli, A. Agarwal, W. Huang, C. N. Parkurst, M. Muratet, K. M. Newberry, S. Meadows, A. Greenfield, Y. Yang, P. Jain, F. K. Kirigin, C. Birchmeier, E. F. Wagner, K. M. Murphy, R. M. Myers, R. Bonneau, and D. R. Littman, “A validated regulatory network for Th17 cell specification,” Cell, vol. 151, no. 2, pp. 289–303, 2012.

[125] D. Marbach, S. Roy, F. Ay, P. E. Meyer, R. Candeias, T. Kahveci, C. A. Bristow, and M. Kellis, “Predictive regulatory models in Drosophila melanogaster by integrative inference of transcriptional networks,” Genome Research,vol. 22, pp. 1334–49, Jul 2012.

[126] A. Pratapa, A. P. Jalihal, J. N. Law, A. Bharadwaj, and T. M. Murali, “Benchmarking algorithms for gene regulatory network inference from single-cell transcriptomic data,” Nature Methods,pp. 1–8, Jan. 2020.

[127] C. Weinreb, S. Wolock, B. K. Tusi, M. Socolovsky, and A. M. Klein, “Fundamental limits on dynamic inference from single-cell snapshots,” Proceedings of the National Academy of Sciences,vol. 115, no. 10, pp. E2467–E2476, 2018.

[128] A. Gioele, L. Manno, R. Soldatov, H. Hochgerner, A. Zeisel, Z. Liu, D. V. Bruggen, J. Guo, E. Sundsträm, G. Castelo-branco, P. Cramer, I. Adameyko, and S. Linnarsson, “RNA velocity in single cells,” Nature,vol. 560, no. 7719, pp. 494–8, 2018.

[129] S. Chen and J. C. Mar, “Evaluating methods of inferring gene regulatory networks highlights their lack of performance for single cell gene expression data,” BMC Bioinformatics,vol. 19, no. 1, p. 232, 2018.

[130] P. Dibaeinia and S. Sinha, “SERGIO: A single-cell expression simulator guided by gene regulatory network,” Cell Systems,vol. 11, pp. 252–271.e11, Sept. 2020.

[131] D. Szklarczyk, A. Franceschini, S. Wyder, K. Forslund, D. Heller, J. Huerta-Cepas, M. Simonovic, A. Roth, A. Santos, K. P. Tsafou, Tsafou, M. Kuhn, P. Bork, L. J. Jensen, and C. von Mering, “STRING v10: protein-protein interaction networks, integrated over the tree of life,” Nucleic Acids Research,vol. 43, pp. D447–D452, Oct 2014.

[132] A. Gitter, Z. Siegfried, M. Klutstein, O. Fornes, B. Oliva, I. Simon, and Z. Bar-Joseph, “Backup in gene regulatory networks explains differences between binding and knockout results,” Molecular Systems Biology, vol. 5, no. 1, 2009.

[133] M. Schrynemackers, R. Kueffner, and P. Geurts, “On protocols and measures for the validation of supervised methods for the inference of biological networks,” Frontiers in Genetics,vol. 4, p. 262, 2013.

[134] D. M. Castro, N. R. d. Veaux, E. R. Miraldi, and R. Bonneau, “Multi-study inference of regulatory networks for more accurate models of gene regulation,” PLOS Computational Biology,vol. 15, p. e1006591, Jan. 2019.

[135] P. Nguyen and R. Braun, “Time-lagged Ordered Lasso for network inference,” BMC Bioinformatics,vol. 19, p. 545, Dec 2018.

[136] M. Yuan and Y. Lin, “Model selection and estimation in regression with grouped variables,” Journal of the Royal Statistical Society. Series B (Statistical Methodology),vol. 68, no. 1, pp. 49–67, 2006.

[137] A. C. Lozano, N. Abe, Y. Liu, and S. Rosset, “Grouped graphical Granger modeling for gene expression regulatory networks discovery,” Bioinformatics,vol. 25, no. 12, pp. i110–i118, 2009.

